# Transcriptional and post-transcriptional regulation and transcriptional memory of chromatin regulators in response to low temperature

**DOI:** 10.1101/757179

**Authors:** K. Vyse, L. Faivre, M. Romich, M. Pagter, D. Schubert, D.K. Hincha, E. Zuther

## Abstract

Chromatin regulation ensures stable repression of stress-inducible genes under non-stress conditions and transcriptional activation and memory of such an activation of those genes when plants are exposed to stress. However, there is only limited knowledge on how chromatin genes are regulated at the transcriptional and post-transcriptional level upon stress exposure and relief from stress. We have therefore set-up a RT-qPCR-based platform for high-throughput transcriptional profiling of a large set of chromatin genes. We find that the expression of a large fraction of these genes is regulated by cold. In addition, we reveal an induction of several DNA and histone demethylase genes and certain histone variants after plants have been shifted back to ambient temperature (deacclimation), suggesting a role in the memory of cold acclimation. We also re-analyse large scale transcriptomic datasets for transcriptional regulation and alternative splicing (AS) of chromatin genes, uncovering an unexpected level of regulation of these genes, particularly at the splicing level. This includes several vernalization regulating genes whose AS results in cold-regulated protein diversity. Overall, we provide a profiling platform for the analysis of chromatin regulatory genes and integrative analyses of their regulation, suggesting a dynamic regulation of key chromatin genes in response to low temperature stress.

## Introduction

Plants are exposed to a multitude of abiotic and biotic stresses during their lifetime and have evolved efficient mechanisms to cope with such events. Stress alleviation relies on numerous changes at the biochemical, physiological and molecular level, largely coordinated by a massive and fast reprogramming of the transcriptome. While it is known that the exposition to chilling temperatures results in the up- and down-regulation of thousands of genes, as well as extensive post-transcriptional regulation of genes such as alternative splicing, all within minutes of cold exposure (Calixto et al., 2018), the precise mechanisms leading to this reprogramming have not been completely elucidated yet. Changes in the transcriptional activity of cold-stress responsive genes upon exposure to low temperatures might be partly achieved through remodeling of the chromatin, rendering it more or less accessible for the transcriptional machinery, thereby affecting the expression levels of the genes. After cold stress ends and deacclimation begins, most stress responsive genes return quickly back to their initial transcriptional levels (Byun et al., 2014;Pagter et al., 2017), suggesting that cold-induced changes to the chromatin might be reversed during deacclimation. Cold induced changes to the chromatin have in particular been studied in the context of vernalization. Vernalization is defined as the acquisition of the competence to flower after prolonged cold treatment, allowing vernalization-responsive plants to flower in spring, under favorable temperature conditions and the appropriate photoperiod. This process relies on epigenetic mechanisms, as cold induces a mitotically stable switch inhibiting the expression of the floral repressor *FLOWERING LOCUS C (FLC)*. The repression of *FLC* is achieved through the action of regulators of the Polycomb-group (Pc-G), which deposit and maintain the repressive tri-methylation of lysine 27 in histone H3 (H3K27me3) upon cold exposure (Song et al., 2012). Changes in the chromatin state were also previously described for cold-inducible genes not involved in vernalization, suggesting the involvement of dynamic chromatin regulation in the induction and repression of cold stress-responsive genes (Kwon et al., 2009;Park et al., 2018). Hyper-acetylation of histone H3K9 in promoter regions of *DREB1*, a main regulator of cold response, was observed in rice and was decreased again when plants were returned to control temperatures, suggesting that deacetylation keeps the gene in an off-state in the absence of cold (Roy et al., 2014). Recently, epigenetic changes involved in cold response were reviewed by different authors (Baulcombe and Dean, 2014;Kim et al., 2015;Asensi-Fabado et al., 2017;Banerjee et al., 2017;Luo et al., 2017;Friedrich et al., 2019). Furthermore, epigenetic changes that establish environmental memory in plants have been described (He and Li, 2018). The existence of memory of a cold priming event, including transcriptional memory, resulting in improved freezing tolerance after a subsequent triggering cold treatment was recently shown (Zuther et al., 2019).

In general, chromatin can be distinguished into euchromatin, consisting of mostly active genes and closed inactive heterochromatin, with a preference for repetitive elements (Allis and Jenuwein, 2016). Chromatin is composed of basic repeating units called nucleosomes, consisting of 145 to 147 bp of DNA wrapped around a histone octamer (Luger et al., 1997). The octamer is formed by core histones H2A, H2B, H3 and H4 and the repeating nucleosomal structure is further linked and stabilized by the linker histone H1 (Luger et al., 1997). This results in the arrangement of higher-order helical structures (Widom, 1989). The nucleosome not only helps in the packaging of the DNA, but is also the primary determinant of DNA accessibility (Luger et al., 1997). Histones, particularly their tails, are extensively post-translationally modified and can undergo a variety of covalent modifications such as acetylation, methylation, phosphorylation and ubiquitination (Zhang and Reinberg, 2001). Histone acetylation is set by histone acetyltransferases (HAT), which transfer the acetyl group of acetyl Coenzyme A to the ε-amino group of lysine side chains (Bannister and Kouzarides, 2011). Histone acetylation is associated with a function as a transcriptional coactivator by neutralizing the positive charge of the lysine. The modification is reversible and the acetyl group can be removed by histone deacetylases to restore the positive charge of the lysine, thus stabilizing the local chromatin architecture. Histone methylation predominantly occurs on the amino acids lysine and arginine. Unlike acetylation, the charge of the histone is not affected (Bannister and Kouzarides, 2011). Lysine residues can be mono-, di- or trimethylated, while arginine can contain one or two methyl groups on its guanidinyl group (Ng et al., 2009). Histone methylation can lead to an active or repressive chromatin state, depending on the modified residue and the number of added methyl groups. Trithorax group (Trx-G) factors are responsible for the deposition of activating methylations on lysine 4 and 36 of histone 3 (H3K4me3 and H3K36me3, respectively), leading to the transcriptional activation of their target genes (del Prete et al., 2015). Their action is antagonized by the proteins of the Polycomb Repressive Complex 2 (PRC2), which mediates H3K27me3. This highly conserved methyltransferase complex contains three orthologues of Enhancer of zeste (E(z)) (CURLY LEAF (CLF), SWINGER (SWN) and MEDEA (MEA)), three orthologues of Suppressor of zeste12 (Su(z)12) (EMBRYONIC FLOWER2 (EMF2), VERNALISATION2 (VRN2) and FERTILISATION INDEPENDENT SEED2 (FIS2)), five orthologues of Multicopy Suppressor of Ira (MSI1-5) and a single copy of Extra Sex Combs (ESC) (FERTILISATION INDEPENDENT ENDOSPERM (FIE) in *Arabidopsis thaliana* (Kleinmanns and Schubert, 2014). Methylation can be reversed by two different classes of histone demethylases: while LSD1-type demethylases can remove one of two methyl groups, JUMONJI-type histone demethylases can counteract mono-, di- or trimethylation (Shi et al., 2004). The second highly conserved Polycomb-Repressive Complex, PRC1, is a histone ubiquitination complex and monoubiquitylates histone H2A (Kleinmanns and Schubert, 2014).

In addition to histone modifications, the state of chromatin can also be affected by DNA methylation, which was found to be linked to gene repression (Hotchkiss, 1948;Razin and Riggs, 1980). DNA methylation (5-methyl cytosine) can be a heritable epigenetic mark (Jin et al., 2011) and is set by DNA methyltransferases. In plants, cytosine can be methylated symmetrically (CG and CHG methylation (where H is any base except G) as well as asymmetrically (CHH)) and these modifications are predominantly found on transposons and other repetitive DNA elements (Zhang et al., 2006). Small RNAs generated by RNA interference (RNAi) target genomic DNA sequences for cytosine methylation (RNA-directed DNA methylation) (Law and Jacobsen, 2010). DNA methylation is reversible and the methyl groups can be removed by demethylases (Penterman et al., 2007). In general, DNA methylation is associated with the repressive state of chromatin, as it alters the accessibility of promoters for transcription factors (Carey et al., 2011).

Dynamic regulation of chromatin is not only achieved by enzymes setting and removing chemical modifications at DNA or histones, but also by replacement of the canonical histones by histone variants, resulting in an immediate loss of histone modifications and a resetting of epigenetic changes (Spiker, 1982). Although nucleosomes are energetically stable, histones can be turned over. Histones H2A and H2B can be exchanged much faster than histones H3 and H4 (Weber and Henikoff, 2014). In *Arabidopsis thaliana*, 13 H2A variants (labelled HTA1-13) exist, and are clustered in 4 groups: H2A, H2A.X, H2A.W and H2A.Z (Kawashima et al., 2015). Histone H3 exists in 15 variants (labelled HTR1-15) distributed in categories including H3.1, H3.3 and CenH3 (Stroud et al., 2012). Different histone variants carry different functions and are located at different parts of the gene to convey regulation. The H3.3 variants are predominantly located towards the 3’-ends of genes and are generally associated with elevation of gene expression, whereas H2A.W binds to heterochromatin with its C-terminal motif KSPKKA and promotes heterochromatin condensation (Yelagandula et al., 2014;Kawashima et al., 2015). Additionally studies have identified H4 variants, however, protein variants have not been described in *A. thaliana* (Kawashima et al., 2015). Lastly, three copies of linker histone H1 exist in *A. thaliana*, H1.1, H1.2 and H1.3 (Kotliński et al., 2016). H1.3 is a stress-inducible histone variant and might be responsible for regulating dynamic DNA methylation (Rutowicz et al., 2015).

While there is ample evidence for a role of chromatin remodeling in the regulation of gene expression in response to cold, relatively little is known about the involvement of specific chromatin regulators. The transcriptional and post-transcriptional regulation of the expression of most of these chromatin modifier genes both during and after cold exposure remains unexplored as well. In the case of vernalization, *VERNALIZATION INSENSITIVE 3* (*VIN3*) is the only VRN gene known to be induced by cold, however, protein level analyses of Pc-G proteins involved in vernalization suggest post-transcriptional regulation of several genes including *VRN2*, *CLF*, *FIE* and *SWN* (Wood et al., 2006).

Here, we set out to analyze the transcriptional and post-transcriptional regulation of chromatin regulatory genes in response to cold stress and following deacclimation using both publically available datasets and generation of a RT-qPCR platform. We identify a potential role for Pc-G proteins in repressing stress-inducible genes under non-stress conditions and substantial transcriptional and post-transcriptional regulation of chromatin regulatory genes. Interestingly, genes involved in vernalization are largely not transcriptionally regulated under short-term (3 days) cold conditions. However, they may be alternatively spliced, resulting in potentially altered protein sequences. Based on data generated with the RT-qPCR platform, we have identified additional cold-inducible chromatin regulatory genes and genes specifically regulated during deacclimation, including DNA demethylases and histone variants.

## Methods

### Plant material

*A. thaliana* accession Col-0 was sown and grown on soil in a climate chamber with 20°C day-time temperature and 6°C night-time temperature in a 14 h light cycle with a light intensity of 180 µE m^-2^ s^-1^ and a humidity of 60% at day and 70% at night. After one week the plants were moved to a short-day climate chamber with the following conditions: 20°C/16°C day/night, 8 h day length, 180 µE m^-2^ s^-1^, humidity of 60%/75% day/night. The plants were kept under these short-day conditions for a week before pricking (10 plants per 10 cm diameter pot). After pricking, the plants were kept for another seven days under short-day conditions before transfer to long-day conditions for another week. The conditions for long-day were 20°C day and 16°C night temperature with a day length of 16 h at a light intensity of 200 µE m^-2^ s^-1^. These four week old plants were used in cold acclimation and deacclimation experiments. For cold acclimation, plants were moved for three days to a growth chamber with a constant temperature of 4°C and a day length of 16 h with a light intensity of 90 µE m^-2^ s^-1^ and a humidity of 70% to 80% (Zuther et al., 2019). For deacclimation, plants were moved back to previous growth conditions for up to 24 h (Pagter et al., 2017). Plant material of 10 individual replicate plants was harvested from non-acclimated (NA) plants (at 8 am), after three days of cold acclimation (at 8 am) and after 2, 4, 6, 12 and 24 h of deacclimation (Deacc). The material was immediately frozen in liquid nitrogen and stored at −80°C before being ground into a fine powder in a ball mill (Retsch, Haan, Germany).

### Selection of genes of interest for the RT-qPCR platform

We focused our selection on chromatin genes associated with epigenetic changes and selected 135 genes for analyses (see Table 1 for abbreviations and Suppl. Table 1 for primer sequences). These include Pc-G genes and Pc-G associated genes (and their paralogs), Trx-G genes, a selection of histone demethylase genes (putative H3K9 and H3K27 demethylases (JUMONJI-type) and LSD1-like histone demethylases), DNA methyltransferase and demethylase and canonical histone and histone variant genes.

**Table 1:**
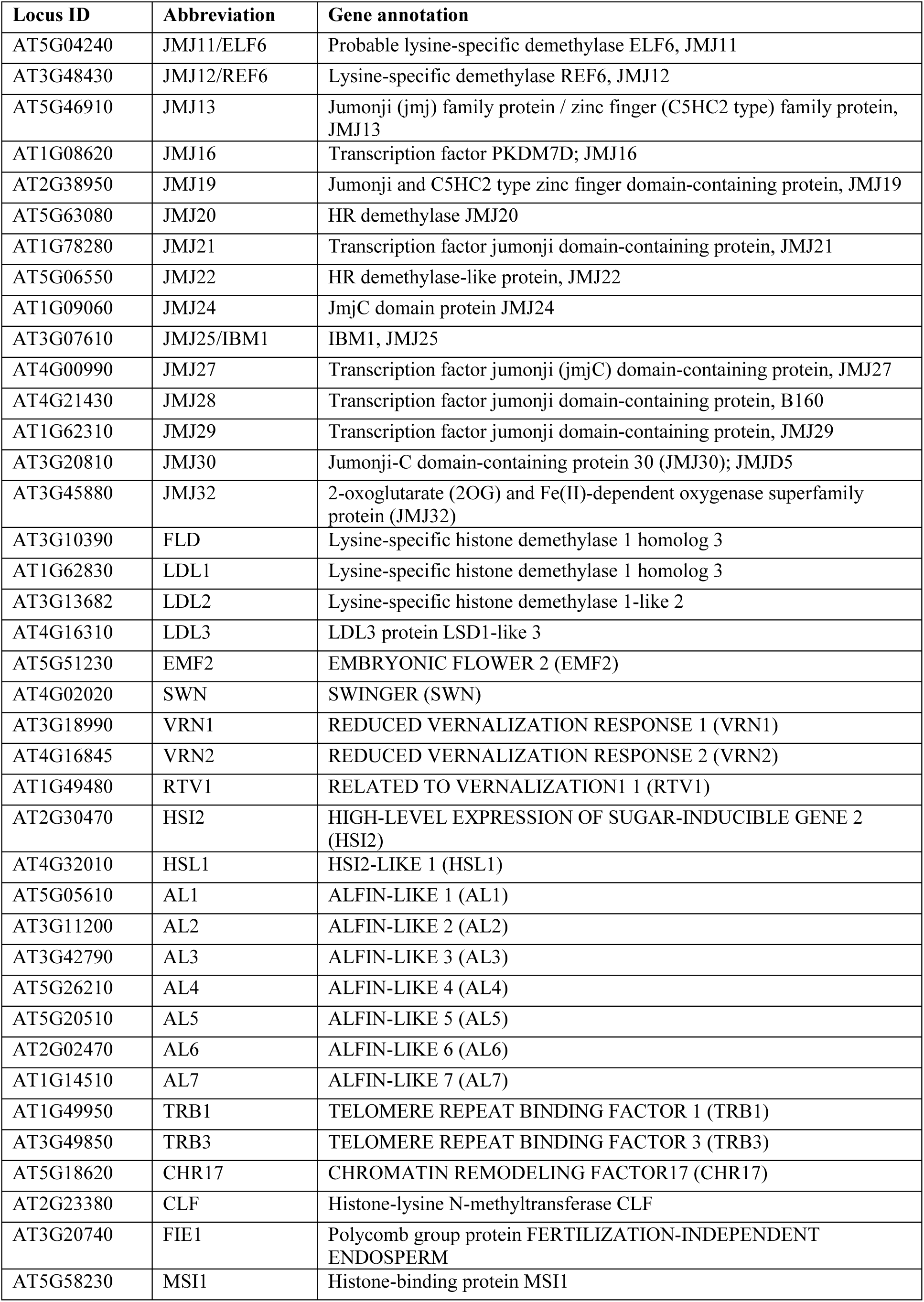
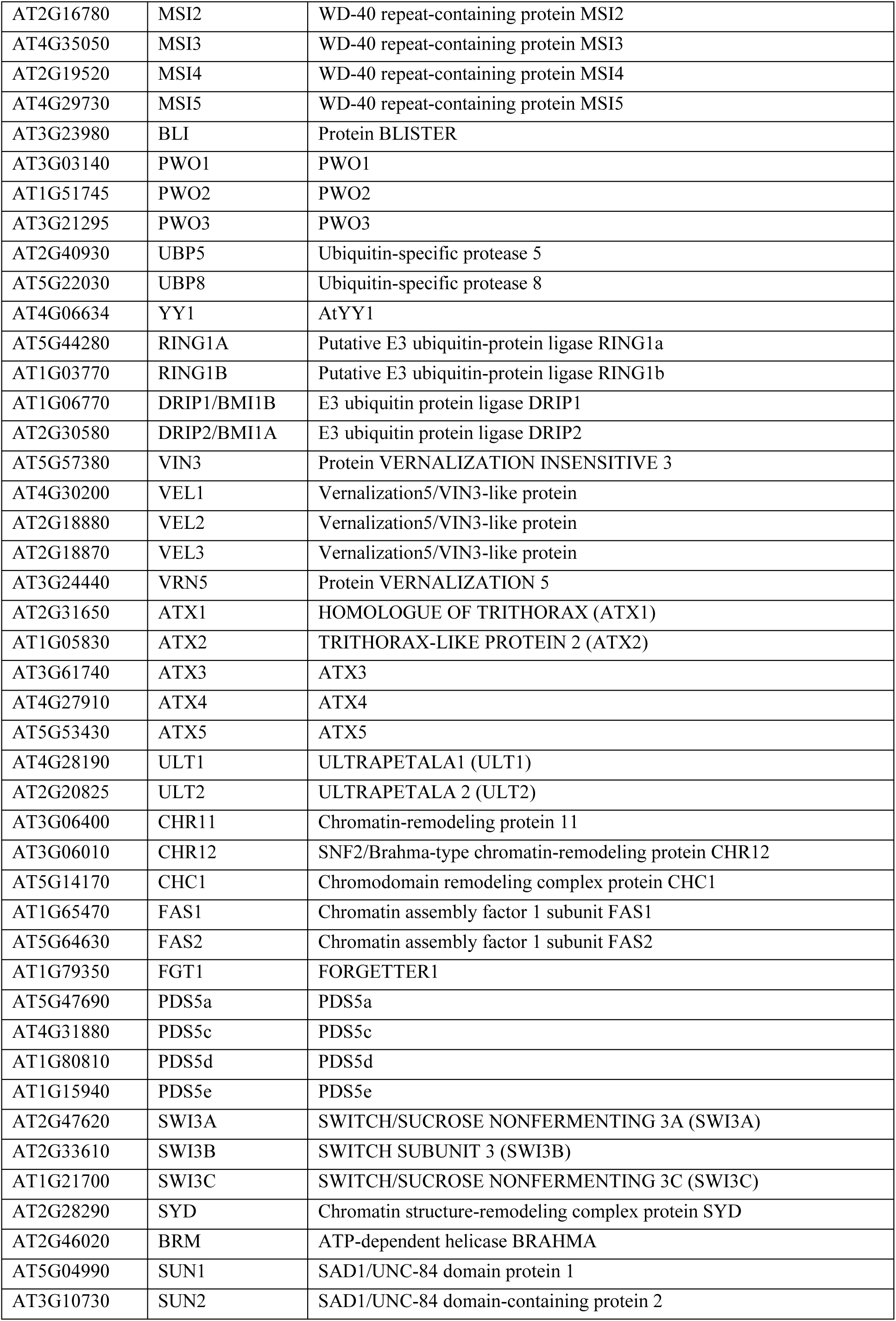
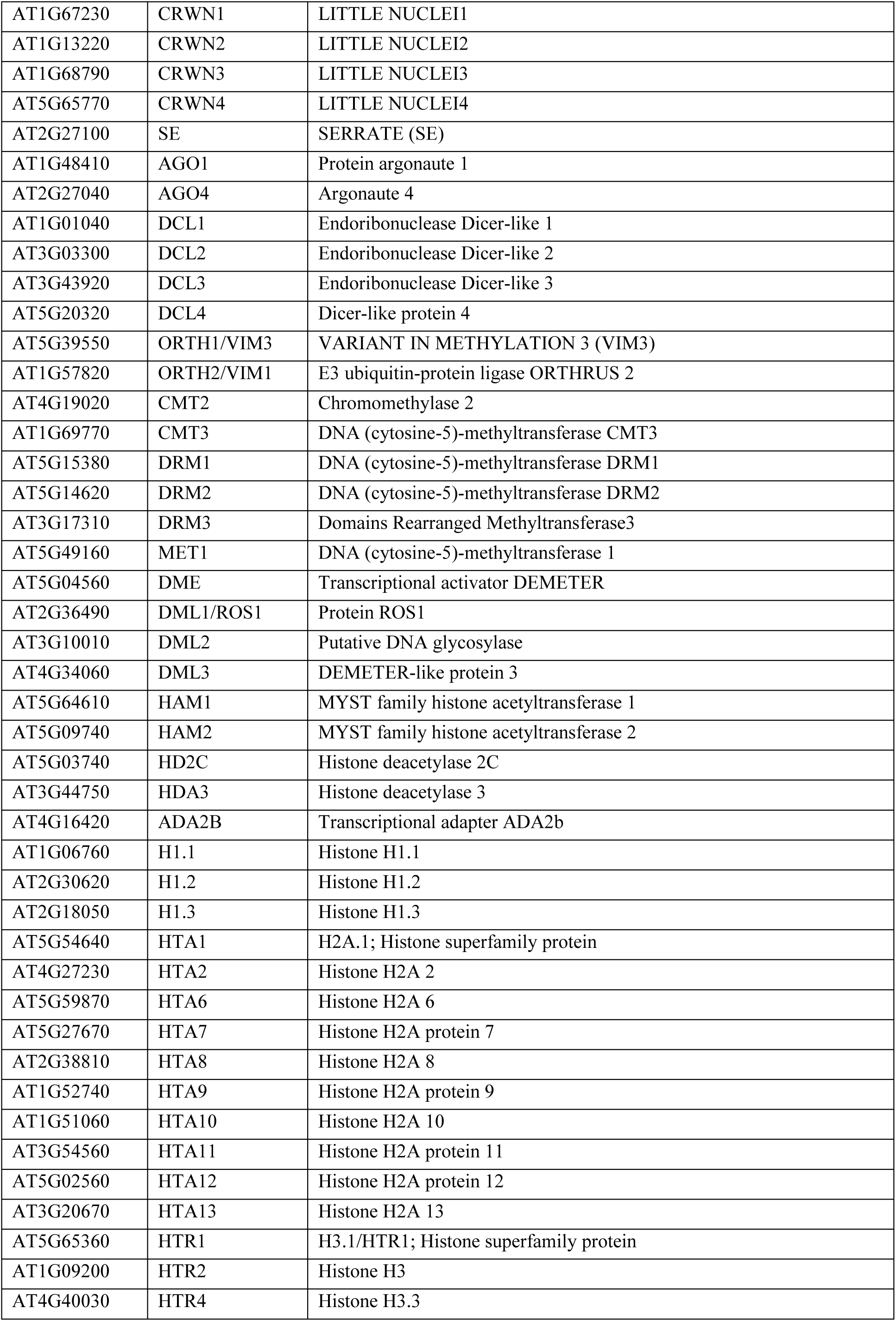
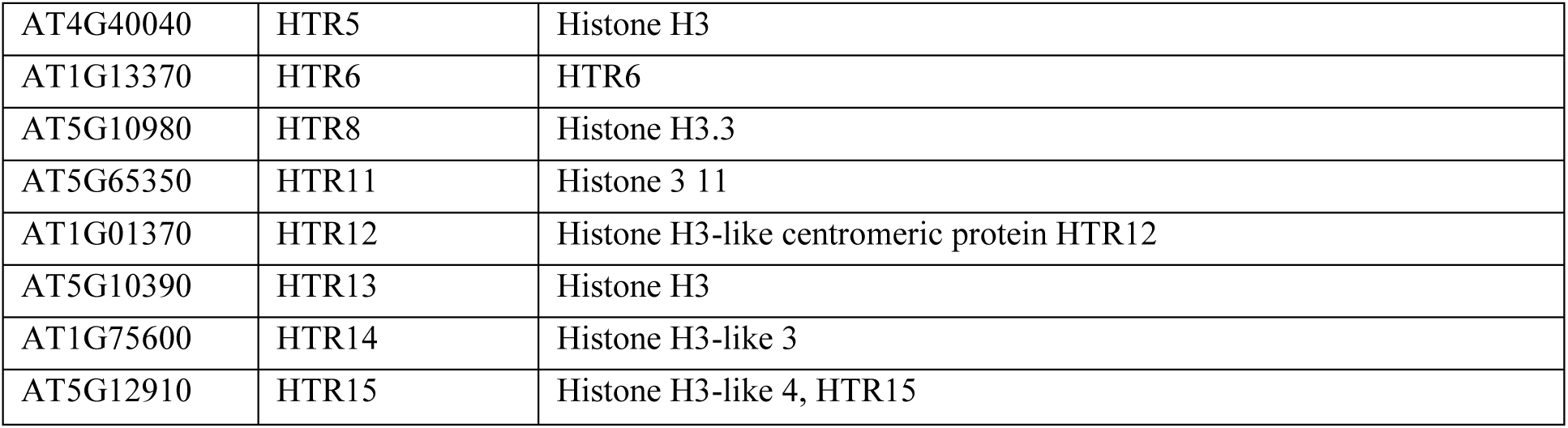
RT-qPCR platform: Nomenclature of 135 genes encoding epigenetic regulators.

### RT-qPCR analysis

RNA was isolated from a pool of five samples consisting of 10 different plants using Trizol reagent (BioSolve BV). RNA was quantified using a NanoDrop-1000 spectrophotometer (Thermo Scientific) and DNA was removed from the samples using a RapidOut DNA Removal Kit (Thermo Scientific). The absence of genomic DNA was tested by RT-qPCR with intron-specific primers (Intron MAF5 AT5G65080) (Zuther et al., 2012). cDNA was synthesised by SuperScript IV Reverse Transcriptase (Invitrogen) and oligo dT18 primers. The quality of the cDNA was tested using primers amplifying the 3’ and 5’ region of *GAPDH* (AT1G13440) (Zuther et al., 2012). cDNA of three independent biological replicates for each time point was used for expression analysis (Pagter et al., 2017).

RT-qPCR was performed for 135 genes of interest (Suppl. Table 1) as previously described (Le et al., 2015). Expression of four housekeeping genes, Actin2 (AT3G18780), EXPRS (AT2g32170), GAPDH (AT1G13440) and PDF (AT1G13320) was measured for each sample on each plate (Suppl. Table 2; compare Zuther et al., 2012). The Ct values were normalised by subtracting the mean of the four housekeeping genes from the Ct value of each gene of interest (ΔCt). Transcript abundance was expressed as 2^-ΔCt^. The log2 fold change of the normalised Ct values was calculated either relative to NA or ACC. Heat maps were constructed in RStudio (R Core Team, 2013;RStudio Team, 2016) using the pheatmap package version 1.0.12.

### Primer design

Primers were either designed in Primer3 or taken from the literature as indicated in Suppl. Table 1. The specifications of the primers were as followed: primer length of 20-24 bases, amplicon size of 60-150 bp, primer melting temperature of 64 ± 3°C, amplicon melting temperature of 75-95°C, G/C content of 45-55%, maximum repetition of a nucleotide of 3 and a G/C clamp of 1. Primers with a binding site near the 3’ end of the respective gene were preferred. For highly homologous genes primer length was increased from 20 to 30 bp.

### Bioinformatic analyses of the overlap of cold-regulated genes and H3K27me3 targets using public data

The dataset of differentially expressed cold-regulated genes was extracted from previously published data (Calixto et al., 2018) Experimental conditions used in this publication were as followed. *Arabidopsis thaliana* Col-0 plants were stratified for four days at 4°C in the dark and then grown in hydroponic culture for five weeks. Growth conditions were 12 h light (150 μE m^-2^ s^-1^)/12 h dark with a constant temperature of 20°C. The 4°C cold treatment was started at dusk. Rosette material was harvested and pooled per sampling point.

H3K27me3 targets were extracted from previously published data (Lafos et al., 2011). In this study H3K27me3 targets genes were identified with whole genome tiling arrays using undifferentiated meristematic cells from the shoot and differentiated leaf tissue from *clavata3* mutant plants (Col-0 background). For this analysis the leaf samples were taken from plants grown for nine weeks under short day conditions (8 h light/16 h dark).

The datasets were compared using the conditional formatting and filter function from Excel.

### Bioinformatic analyses of the cold regulation of chromatin modifier genes using public data

For the investigation of the regulation of chromatin genes during cold exposure, a list of chromatin regulator genes was extracted from the ChromDB database (version of the 30^th^ of July, 2011) (Gendler et al., 2008). This list was overlapped with the lists of genes either differentially expressed or differentially spliced at any time point during cold exposure (Calixto et al., 2018) using the VennDiagram R package.

The analysis of the cold regulation of vernalization actors was performed by overlapping a list of genes involved in vernalization regulation with the lists of cold-regulated and cold-alternatively spliced genes previously described The Venn diagram was plotted using the VennDiagram R package. The expression profiles of genes showing differential splicing upon cold exposure were generated and downloaded using the webservice at https://wyguo.shinyapps.io/atrtd2_profile_app/ (Zhang et al., 2017;Calixto et al., 2018). The translations of the differential usage transcripts were extracted from AtRTD2 (Zhang et al., 2017) and aligned to one another using the Needle algorithm (Madeira et al., 2019) to identify potential differences in amino acid sequences caused by cold exposure. The alignments were visualized using Multiple Align Show from the Sequence Manipulation Suite (Stothard, 2000).

### Statistics

The statistical significance of overlaps between different groups of genes was calculated using http://nemates.org/MA/progs/overlap_stats.html. The significance of gene expression changes was analysed using an unpaired two-sided t-test, performed in RStudio (R Core Team, 2013;RStudio Team, 2016). The significance levels are presented as followed: *, 0.05>p>0.01; **, 0.01>p>0.001; ***, p < 0.001.

## Results

### Enrichment of H3K27me3 target genes in early, but not late cold-inducible genes

Expression of stress-inducible genes needs to be tightly controlled to prevent costly induction of plant defense responses in the absence of abiotic and biotic stresses. We hypothesized that stress-inducible genes are epigenetically silenced under non-stress conditions and therefore analysed the prevalence of the main epigenetic silencing mark H3K27me3 in cold-regulated genes. We compared the H3K27me3 target genes in mature leaves (profiled under NA control conditions) with genes regulated by short-term (3 h and 6 h) or long-term (3 d) cold (Lafos et al., 2011;Calixto et al., 2018) (Fig. 1, Suppl. Table 3). H3K27me3 target genes were significantly enriched among the early cold-inducible genes (induced after 3 h in the cold), but not in later inducible genes (induced only after 6 h or after 3 d). The opposite pattern was revealed for genes down-regulated in the cold: H3K27me3 target genes are underrepresented in the early (transiently) repressed genes, but enriched in the very late repressed genes (3 d).

**Figure 1:**
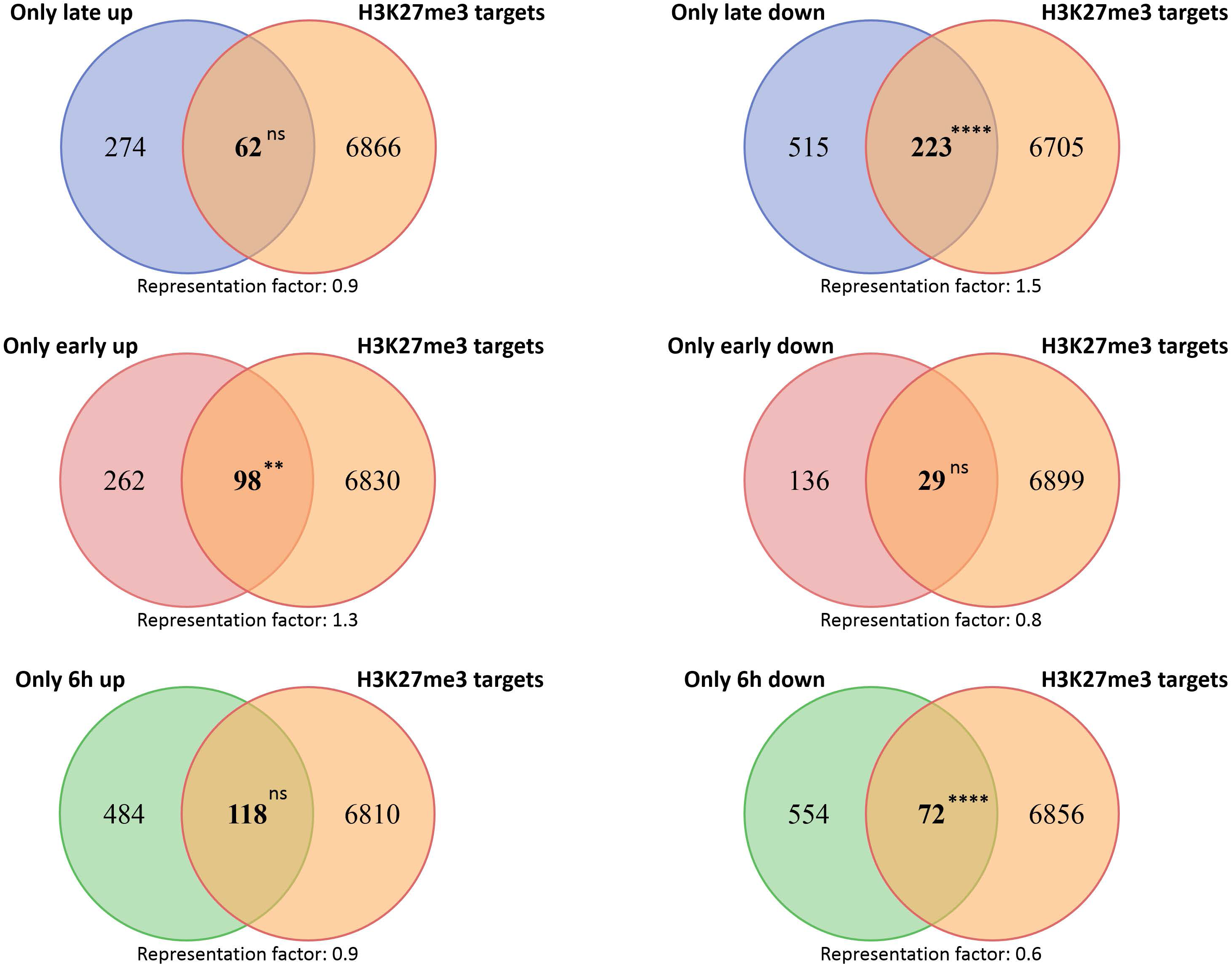
Venn diagrams showing the overlap of cold-regulated genes with H3K27me3 target genes. Asterisks indicate **** = *P* < 0.0001; *** = *P* < 0.001; ** = *P* < 0.01; * = *P* < 0.05. For details see Suppl. Table 3.

### Transcriptional regulation and alternative splicing of chromatin genes during cold exposure

As our bioinformatic analyses suggested resetting of H3K27me3 at many early cold-inducible genes, we wondered whether Pc-G, Pc-G associated and Pc-G antagonist genes and other chromatin genes are subject to transcriptional and/or post-transcriptional regulation upon cold-exposure. We analysed this in two ways: first, we generated a RT-qPCR platform permitting expression analyses of Pc-G, Pc-G associated and Pc-G antagonist genes, histone and histone variant genes and DNA methyltransferases and demethylases (in total 135 genes, see Methods and below). In addition, we extracted chromatin genes from the Chromatin Database ChromDB (Gendler et al., 2008) and analysed their transcriptional regulation and alternative splicing using a published dataset (Calixto et al., 2018). While chromatin regulatory genes were not significantly enriched among the genes differentially expressed in the cold (DE genes), they were highly enriched among the differentially alternatively spliced genes (DAS genes) (Fig. 2). Only 14 out of 511 genes from ChromDB were both differentially expressed and spliced in the cold.

**Figure 2:**
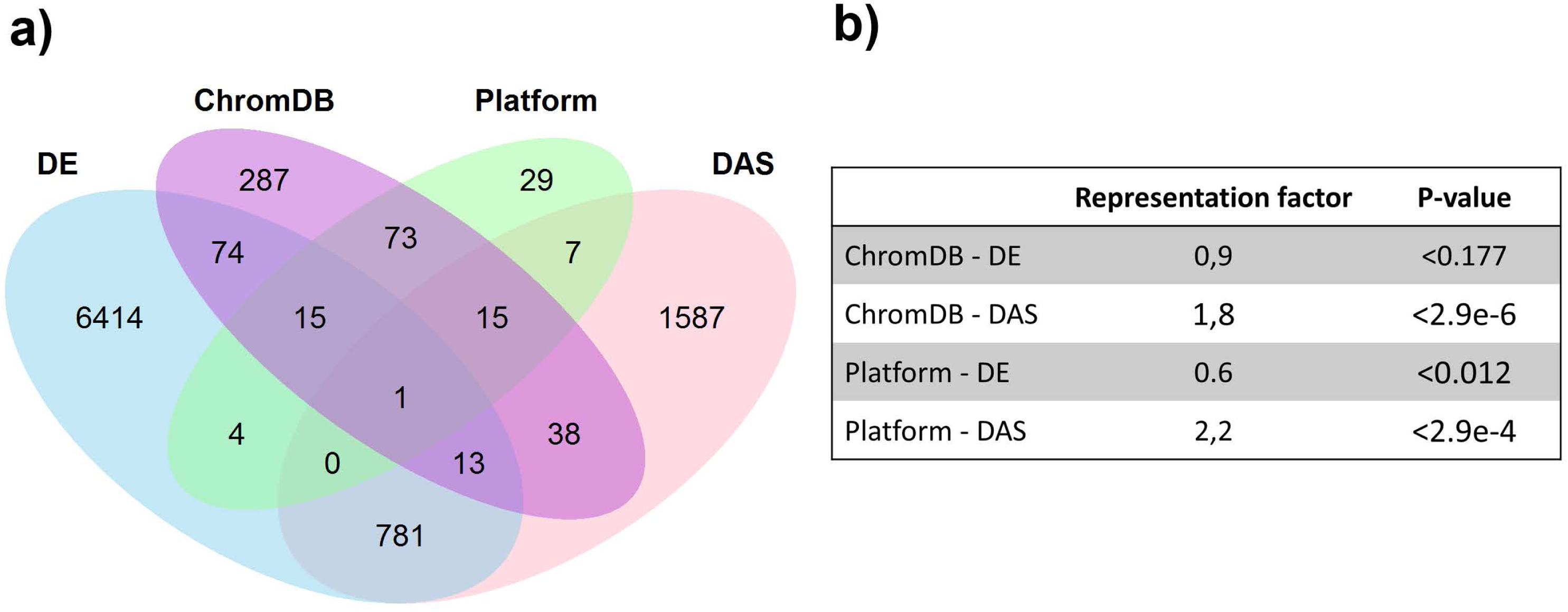
a) Venn diagrams showing the overlap of cold-regulated genes with chromatin regulatory genes. Overlaps were generated for differentially expressed (DE) and differentially alternatively spliced (DAS) genes, based on published data (Calixto et al., 2018), genes extracted from the Chromatin database (ChromDB) and present on the RT-qPCR platform (Platform). b) Significance of overlap and representation factor for the different comparisons. For details see Suppl. Table 4.

While chromatin regulatory genes were not enriched among cold-regulated genes, interesting regulation patterns of several gene families were identified. Among the transcriptionally regulated genes were six histones and histone variants (*HTR2*, *HTR6*, *HTR11*; *HTA6*, *HTA12*; *H1.3*) and all three paralogues of the DNA demethylase *DEMETER*, *DML1*/*ROS1*, *DML2*, *DML3* (Suppl. Table 4). Genes involved in vernalization were not differentially expressed in response to cold, however, several vernalization regulators showed alternative splicing events (seven out of 15 genes) (Table 2). These include *SR45*, *EMF2, VRN2*, *VRN5*, *VEL1*, *SWN* and *HSL1*. For most genes, alternative splicing resulted in repression or induction of alternative variants which encode slightly altered proteins, e.g. for VRN2 a cold-induced alternative transcript translates into a protein with an addition of the amino acids QL at position 304 which is within the highly conserved VEFS domain (Suppl. Fig. 1). Interestingly, among the genes that were both differentially expressed and spliced was the H3K27me3 demethylase *JMJ30*. Its close paralogue, *JMJ32* was also differentially expressed but not alternatively spliced. Overall, our bioinformatic analyses detected a widespread differential expression and splicing of chromatin regulators. Particularly, cold-induced alternative splicing of vernalization regulators is interesting and consistent with previously reported post-transcriptional regulation of *VRN2*, *FIE*, *CLF* and *SWN* (Wood et al., 2006). Whether the proteins generated by alternative splicing exhibit different functional properties or have different interaction partners remains to be discovered.

**Table 2:**
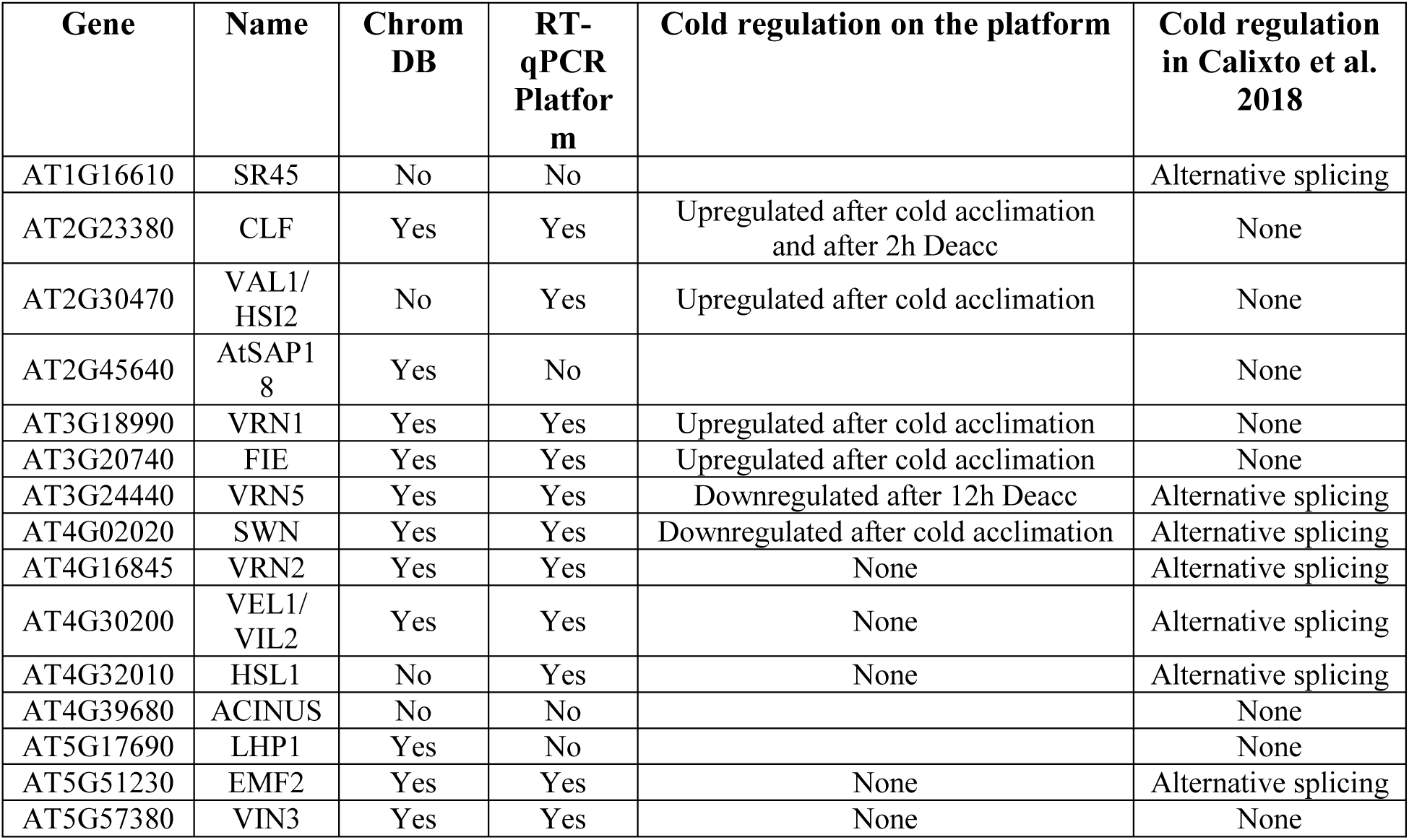
Summary of the regulation of genes involved/related to vernalization. Columns indicate the presence in ChromDB, on the RT-qPCR platform and whether expression regulation has been detected in previously published data (Calixto et al., 2018) and/or in the RT-qPCR platform. Alternative transcripts and encoded proteins are displayed in Suppl. Figure 1.

| Gene | Name | Chrom DB | RT-qPCR Platform | Cold regulation on the platform | Cold regulation in Calixto et al. 2018 |
| --- | --- | --- | --- | --- | --- |
| AT1G16610 | SR45 | No | No |  | Alternative splicing |
| AT2G23380 | CLF | Yes | Yes | Upregulated after cold acclimation and after 2h Deacc | None |
| AT2G30470 | VAL1/HSI2 | No | Yes | Upregulated after cold acclimation | None |
| AT2G45640 | AtSAP18 | Yes | No |  | None |
| AT3G18990 | VRN1 | Yes | Yes | Upregulated after cold acclimation | None |
| AT3G20740 | FIE | Yes | Yes | Upregulated after cold acclimation | None |
| AT3G24440 | VRN5 | Yes | Yes | Downregulated after 12h Deacc | Alternative splicing |
| AT4G02020 | SWN | Yes | Yes | Downregulated after cold acclimation | Alternative splicing |
| AT4G16845 | VRN2 | Yes | Yes | None | Alternative splicing |
| AT4G30200 | VEL1/VIL2 | Yes | Yes | None | Alternative splicing |
| AT4G32010 | HSL1 | No | Yes | None | Alternative splicing |
| AT4G39680 | ACINUS | No | No |  | None |
| AT5G17690 | LHP1 | Yes | No |  | None |
| AT5G51230 | EMF2 | Yes | Yes | None | Alternative splicing |
| AT5G57380 | VIN3 | Yes | Yes | None | None |

### Expression of genes encoding proteins involved in epigenetic processes during cold acclimation and deacclimation

To allow expression analyses of Pc-G, Trx-G and associated genes, histone genes and genes involved in DNA methylation under various conditions, we set up a RT-qPCR platform. The expression of the 135 selected genes was analyzed in samples from non-acclimated (NA), cold-acclimated (ACC) and deacclimated (Deacc) plants after 2 h, 4 h, 6 h, 12 h and 24 h of deacclimation. Due to extremely low expression levels throughout all samples, gene expression data for *JUMONJI* (*jmc*) domain*-containing protein 14* (*JMJ14*) (AT4G20400)*, Maternal affect embryo arrest 27/*(*JMJ15*) (AT2G34880), (*JMJ17*) (At1g63490), (*JMJ18*) (AT1G30810), (*JMJ26*) (At1g11950)*, VP1/ABI3-like 3* (*VAL3*) (AT4G21550)*, Probably E3-ubiquitin protein ligase* (*DRIPH*) (AT3G23060)*, Chromomethylase 1* (*CMT1*) (AT1G80740), and *male-gamete-specific histone H3* (*MGH3*) (AT1G19890) were not further considered, resulting in 135 investigated genes.

Expression changes of these genes during cold acclimation and subsequent deacclimation are shown in five heat maps complied according to the function of the respective proteins in epigenetic regulation. The corresponding normalized 2^-ΔCt^ values are shown in Suppl. Table 5.

The expression of nine out of 19 genes encoding JUMONJI-type and lysine specific 1A-type (LSD1) histone demethylases was significantly upregulated after cold acclimation. Out of these nine genes, *JMJ11/ELF6, JMJ19, JMJ22, JMJ27, JMJ28 and JMJ30* showed the highest log2 fold change compared to NA (Fig. 3). During deacclimation the expression of these genes decreased over time which is additionally illustrated in the comparison of gene expression levels at all deacclimation time points with the expression at ACC conditions (Suppl. Fig. 2, Suppl. Table 6). However, most genes displayed a drop in expression after 4 h Deacc followed by a slight increase at 6 h Deacc. Almost all genes showed a significant downregulation of the expression relative to ACC after 24 h Deacc (Suppl. Fig. 2) and no significant differences compared to NA conditions.

**Figure 3:**
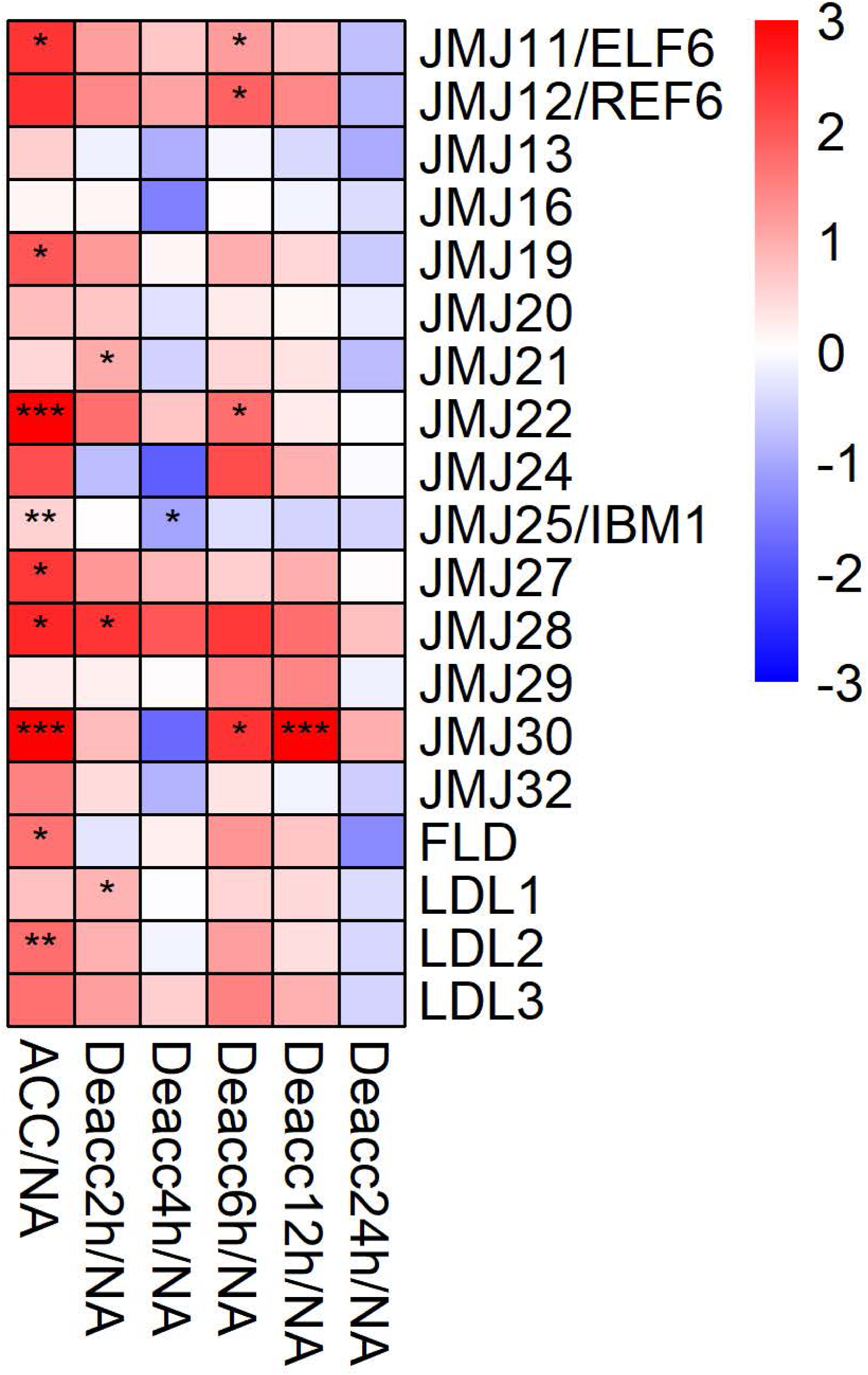
Expression changes of genes encoding JUMONJI-type and LSD1-type histone demethylases after cold acclimation (ACC) and after 2 h, 4 h, 6 h, 12 h and 24 h of deacclimation (Deacc). Gene expression is presented as log2 fold change to non-acclimated conditions (NA) (Suppl. Table 6). The scale of log2 fold changes ranges from −3 (blue) to 3 (red) with a median of 0 (white). Significance levels are indicated relative to NA: ***, *P* < 0.001; **, *P* < 0.01; *, *P* < 0.05 *.

Forty genes encoding members of the Pc-G related protein family were included in the expression analysis by RT-qPCR (Fig. 4, Suppl. Fig. 3). Similarly to the JUMONJI-type and LSD1-type histone demethylase families, most Pc-G related genes displayed an upregulation during cold acclimation, with the highest upregulation for *CLF,* four *WD40 repeat containing proteins MSI1-4* and *VRN1*. Altogether 13 genes encoding Pc-G related proteins were significantly upregulated, while *SWN*, *DRIP2/BMI1A* and *VEL3* exhibited a downregulation at ACC. Most genes with a strong upregulation at ACC kept their higher expression levels over 12 h of deacclimation before they were significantly downregulated in comparison to ACC after 24 h (Suppl. Fig. 3, Suppl. Table 6). After 6 h Deacc their expression was either increased transiently before returning to the initial NA expression levels or was continuously downregulated during the 24 h of deacclimation (Fig. 4). Ten genes displayed a decrease in expression during deacclimation, which was significant in comparison to ACC over several time points, including *AL3, CLF*, *FIE1*, *MSI1-MSI5*, *YY1* and *VIN3* (Suppl. Fig. 3). *VEL2* was the only gene with a significant up-regulation in comparison to ACC after 24 h Deacc (Suppl. Fig. 3). *DRIP2/BMI1A* on the other hand was the only gene of the Pc-G related protein family with a significantly reduced expression at 24 h Deacc compared to NA.

**Figure 4:**
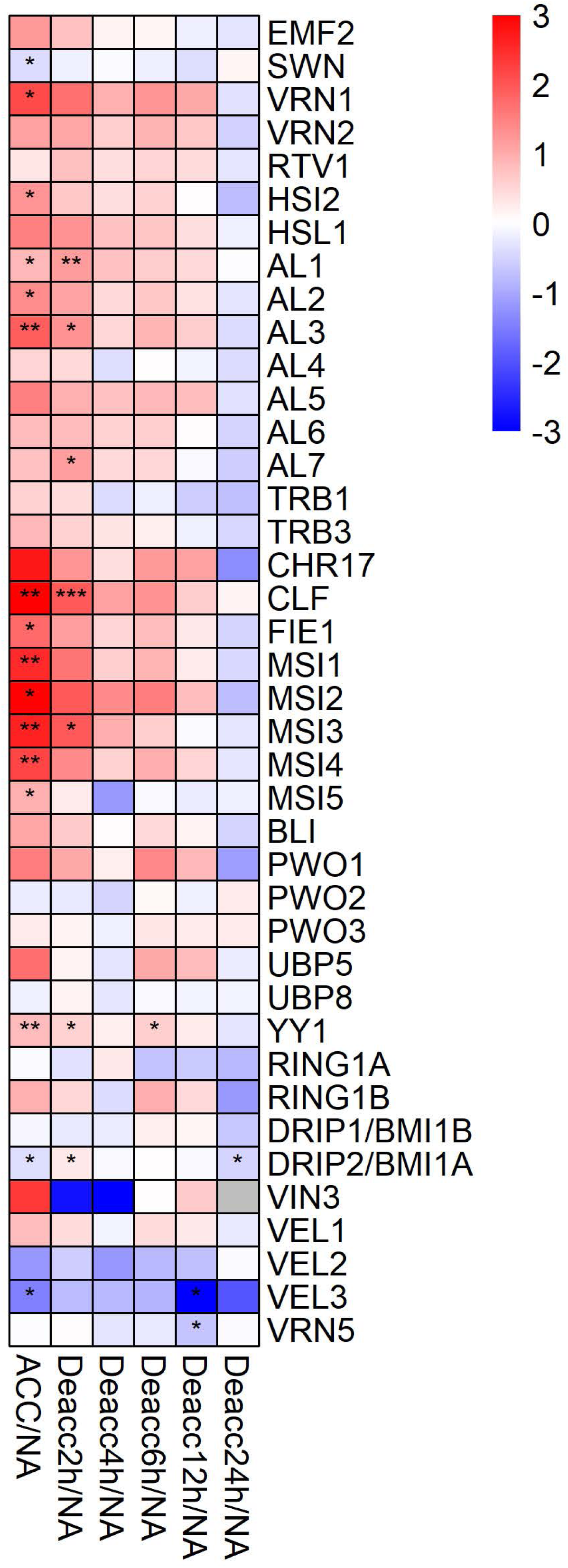
Expression changes of genes encoding proteins of the polycomb group (Pc-G) family after cold acclimation (ACC) and after 2 h, 4 h, 6 h, 12 h and 24 h of deacclimation (Deacc). Gene expression is presented as log2 fold change to non-acclimated conditions (NA) (Suppl. Table 6). The scale of log2 fold changes ranges from −3 (blue) to 3 (red) with a median of 0 (white). Significance levels are indicated relative to NA: ***, *P* < 0.001; **, *P* < 0.01; *, *P* < 0.05 *.

The expression of 22 genes encoding Trithorax-group (Trx-G) related proteins and chromatin remodelers was also investigated (Fig. 5, Suppl. Table 6). Eight of these genes were significantly induced during cold acclimation (*ATX1*, *ATX4*, *ULT2*, *CHR12*, *FAS1*, *FAS2*, *PDS5d* and *SWI3A*). After 2 h, 6 h and 12 h Deacc only two, four and two of these genes, respectively, were still significantly induced with only *SWI3A* showing a stable significant induction over almost all time points. A significant transiently changed expression over two time points during deacclimation compared to NA was only evident for *PDS5e* at 6 h and 12 h. At 24 h Deacc expression changes caused by cold acclimation were mostly reversed and expression of all genes reached similar levels as under NA conditions, except for *ULT1*, which was significantly downregulated. *ATX5* displayed a transient upregulation in comparison to ACC conditions till 12 h Deacc (Suppl. Fig. 4).

**Figure 5:**
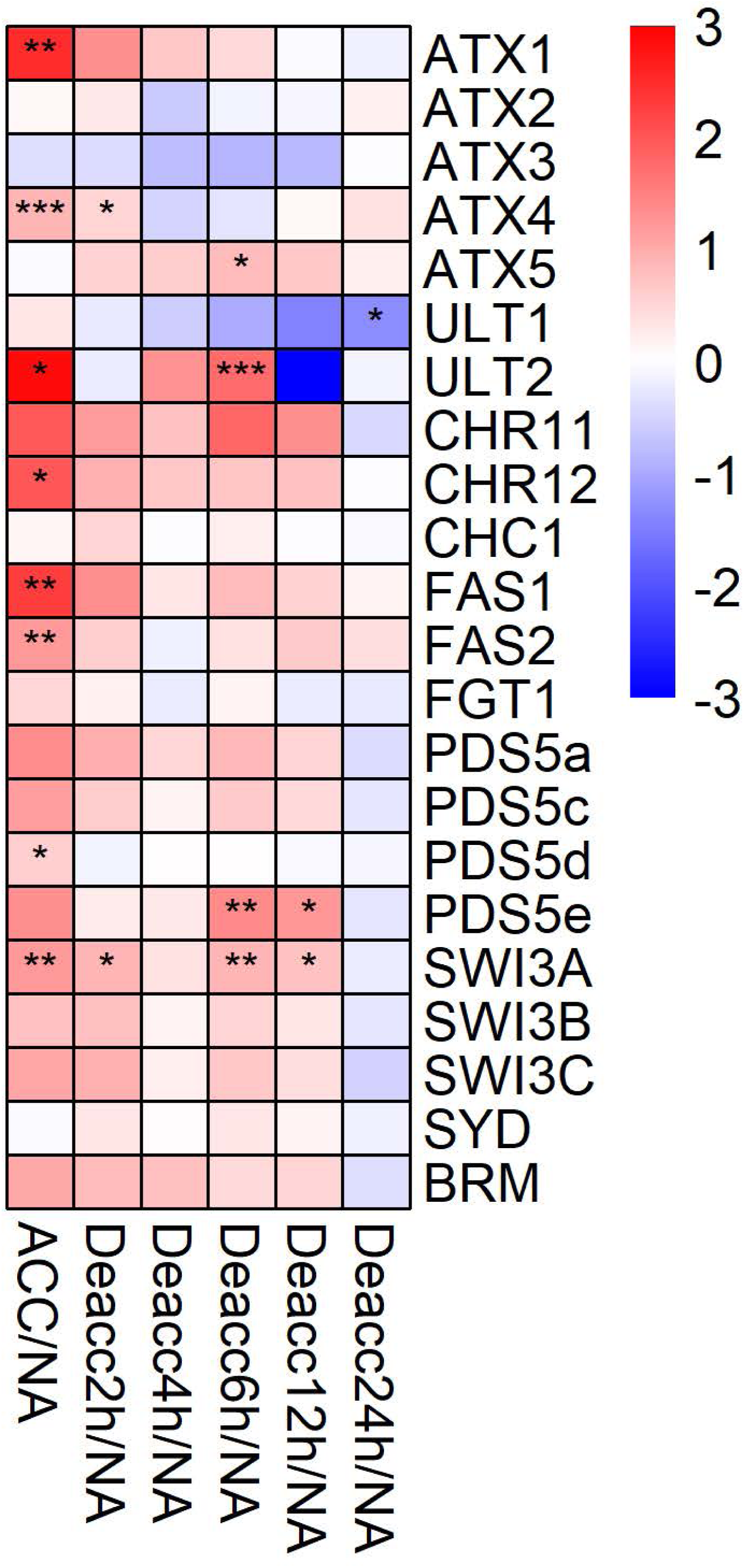
Expression changes of genes encoding proteins of the trithorax group (Trx-G) after cold acclimation (ACC) and after 2 h, 4 h, 6 h, 12 h and 24 h of deacclimation (Deacc). Gene expression is presented as log2 fold change to non-acclimated conditions (NA) (Suppl. Table 6). The scale of log2 fold changes ranges from −3 (blue) to 3 (red) with a median of 0 (white). Significance levels are indicated relative to NA: ***, *P* < 0.001; **, *P* < 0.01; *, *P* < 0.05 *.

Further, the expression of 25 genes encoding proteins acting in chromosome-nuclear envelope (Chr-NE) interactions, RNA interference and DNA methylation was measured (Fig. 6, Suppl. Fig 5). Five genes of this group were significantly differentially expressed at ACC compared to NA (Fig. 6), *SE*, *DCL1*, *ORTH1/VIM3*, *MET1* and *DML3*. Only *DML3* was significantly reduced in its expression under ACC compared to NA conditions. For seven genes of this group expression decreased significantly at different time points of deacclimation compared to ACC, *SUN2*, *SE*, *DCL1*, *ORTH1/VIM3*, *DRM3*, *MET1* and *DML3* (Suppl. Fig. 5, Suppl. Table 6). In contrast, especially *CMT2*, *DRM1* and *DME* displayed a stable upregulation until 12 h Deacc or 6 h Deacc (Fig. 6). *DML3* was highly induced at almost all time points of deacclimation compared to ACC and was together with *DRM3* and *DML1/ROS1* still significantly upregulated compared to ACC at 24 h Deacc (Suppl. Fig. 5). Lastly, changes in expression levels of 29 genes that encode histone acetyltransferases (HAC), deacetylases (HDAC) or histone variants were investigated (Fig. 7, Suppl. Fig. 6). Fourteen of the selected genes were significantly up-regulated after three days of cold acclimation, including *HDA3*, *HTR1* and *HTR2*, *HTR12* and *HTR13,* which showed the highest induction (Suppl. Table 6). Strikingly, this up-regulation was still present after 2 h Deacc for 12 of these genes and became significant for three additional ones, *HTA1*, *HTA10* and *HTR5*. About half of the genes displayed up-regulated expression in comparison to NA over 6 h of deacclimation before they were downregulated after 24 h. Especially for *H1.1, HTR1* and *HTR5*, a more stable upregulation was observed which was still significant after 12 h Deacc. These genes displayed an increase in expression after cold acclimation, followed by a slight decrease up to 4 h of deacclimation compared to NA conditions. From 4 h to 12 h Deacc, expression increased again before returning to the NA level after 24 h, similarly to the pattern in most JUMONJI family genes.

**Figure 6:**
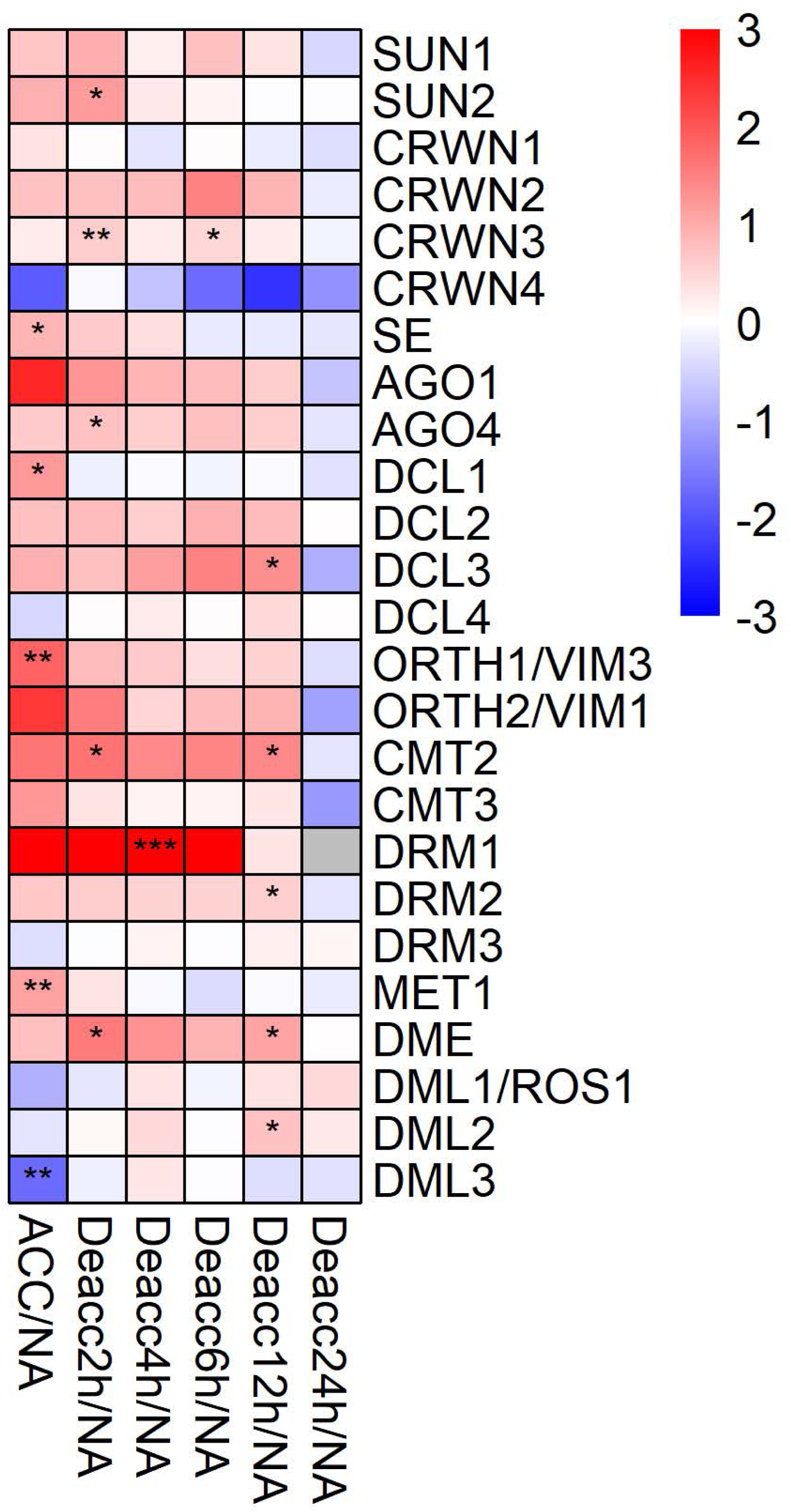
Expression changes of genes encoding proteins acting in chromosome-nuclear envelope (Chr-NE) interactions, RNA interference and methylation after cold acclimation (ACC) and after 2 h, 4 h, 6 h, 12 h and 24 h of deacclimation (Deacc). Gene expression is presented as log2 fold change to non-acclimated conditions (NA) (Suppl. Table 6). The scale of log2 fold changes ranges from −3 (blue) to 3 (red) with a median of 0 (white). Significance levels are indicated relative to NA: ***, *P* < 0.001; **, *P* < 0.01; *, *P* < 0.05 *.

**Figure 7:**
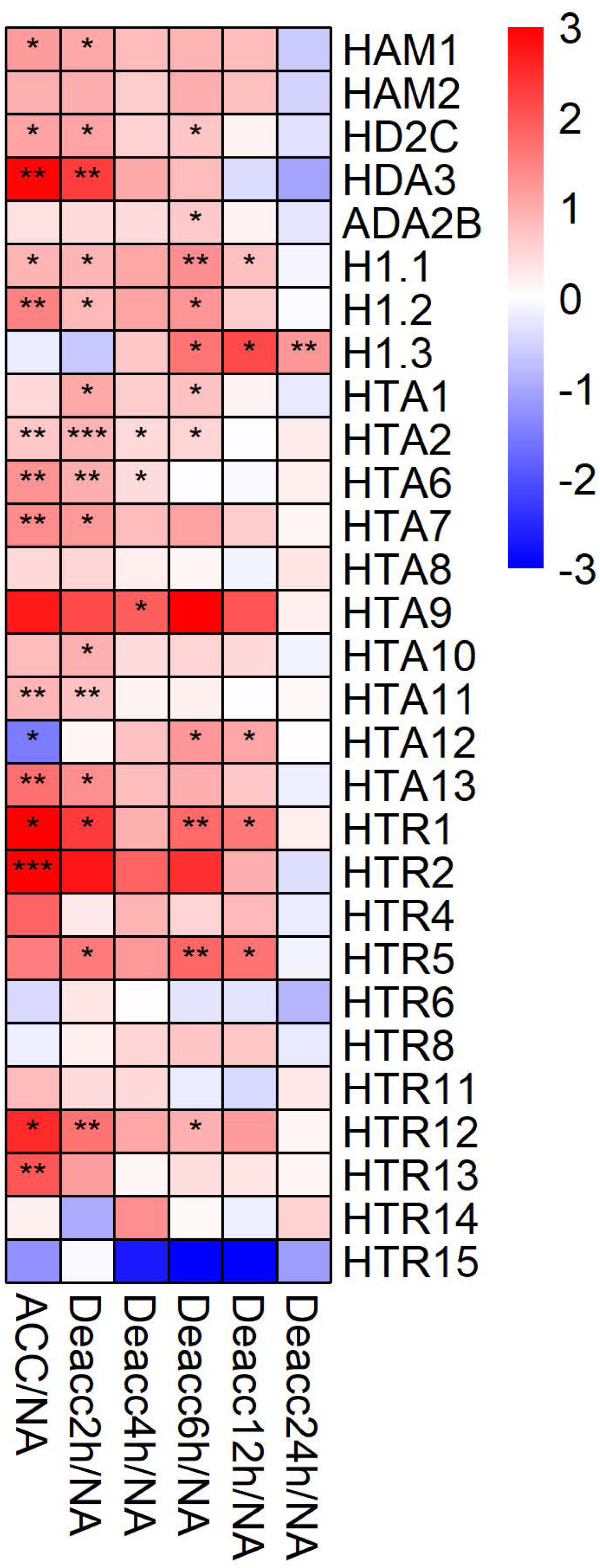
Expression changes of genes encoding histone acetyltransferases (HAC), deacetylases (HDAC) or histone variants after cold acclimation (ACC) and after 2 h, 4 h, 6 h, 12 h and 24 h of deacclimation (Deacc). Gene expression is presented as log2 fold change to non-acclimated conditions (NA) (Suppl. Table 6). The scale of log2 fold changes ranges from −3 (blue) to 3 (red) with a median of 0 (white). Significance levels are indicated relative to NA: ***, *P* < 0.001; **, *P* < 0.01; *, *P* < 0.05 *.

Interestingly, *HTA12* was upregulated at later time points of deacclimation (6 h and 12 h Deacc) after exhibiting the largest decrease in expression after cold acclimation in comparison to NA. Consistently, this gene was significantly induced compared to ACC along the whole deacclimation time course (Suppl. Fig. 6). Lastly, *H1.3* was the only investigated gene that displayed a decrease after cold acclimation and an increase throughout the 24 h Deacc (Suppl. Fig. 6, Suppl. Table 6).

## Discussion

### H3K27me3 preferentially targets early inducible and late repressed genes

Chromatin and chromatin modifications contribute to the regulation of stress-regulated genes at various layers: (1) the repression of stress-inducible genes in non-stress conditions (by repressive chromatin), (2) the activation or repression of genes immediately after stress exposure (by the acquisition of active or repressive chromatin, respectively), (3) the sustained activation or repression in non-stress conditions after exposure to the stress and (4) the transcriptional memory of a stress (in non-stress conditions), permitting primed gene regulation when exposed again to the stress (Friedrich et al., 2019). It is therefore conceivable that chromatin genes and the activity of their gene products are regulated at various layers during different phases of stress exposure and relief. Pc-G mediated H3K27me3 is one of the key repressive chromatin modifications and targets thousands of genes in non-stress conditions, which are developmental regulators, tissue-specifically regulated genes and stress-responsive genes (Lafos et al., 2011). By bioinformatics comparison of H3K27me3 target genes and a detailed kinetic analysis of cold-regulated genes, we revealed an enrichment of H3K27me3 target genes among the early inducible genes. As the early inducible genes are required to trigger a cascade of gene regulatory networks to permit cold acclimation, it is likely particularly important to control their tight repression in non-stress conditions. Resetting of silencing (and possibly of H3K27me3) may occur via different mechanisms, the enzymatic removal by histone demethylases, the addition of active modifications leading to a bivalent chromatin state, the exchange of histones by histone variants and the removal or sliding of nucleosomes by chromatin remodeling complexes. Interestingly, several genes potentially regulating dynamic changes in H3K27me3 are induced by cold, including the H3K27me3 demethylase JMJ30 (Gan et al., 2014), the Trx-G proteins ATX1 (a H3K4 methyltransferase) and ULT1 (Alvarez-Venegas et al., 2003;Carles and Fletcher, 2009) and diverse histone variants.

Our analyses also revealed that H3K27me3 target genes (identified in warm conditions) are overrepresented among the late repressed genes in the cold (Fig. 1). As the changes in H3K27me3 occupancy upon cold exposure have not yet been revealed it is unclear whether the late repressed H3K27me3 target genes acquire a higher level of H3K27me3 or H3K27me3 in more tissues in the cold (compared to warm conditions). One such example is *FLOWERING LOCUS C* (*FLC*) which carries H3K27me3 in the warmth but a higher level of H3K27me3 upon cold exposure/vernalization (Schubert et al., 2006). These genes may be marked for long-term repression in the cold, but further chromatin analyses will be required to elucidate this.

### Expression analysis of genes mediating epigenetic changes

In general, our analysis indicated that cold acclimation had a strong influence on the expression of the selected epigenetics-related genes. This is in agreement with the finding that cold stress enhanced the accessibility of chromatin and bivalent histone modifications of active genes in potato (Zeng et al., 2019). After 24 h of deacclimation, expression of the investigated genes had largely returned to the NA status. This is in agreement to our earlier data on the expression of genes encoding transcription factors, investigated by RT-qPCR, and global gene expression, investigated by microarray hybridization (Pagter et al., 2017).

### Genes encoding JUMONJI-type and LSD1 histone demethylases are upregulated after cold acclimation

JUMONJI-type and LSD1-type enzymes are histone demethylases. For several years histone methylation was thought to be irreversible until the discovery of human LSD1 (Mosammaparast and Shi, 2010). Although members of both enzyme families target histone methylation, they have different structures as well as targets. The JUMONJI family is defined by a JUMONJI C (JmjC) domain consisting of two histidines and one glutamate residue to chelate the catalytic iron, which is essential for its function (Mosammaparast and Shi, 2010). JmjC proteins are able to demethylate tri-methylated lysines, including those on H3K9, H3K27 and H3K36 (Mosammaparast and Shi, 2010;Gan et al., 2014). JmjC-domain proteins further reverse trimethylated H3K4 to its mono- or dimethylated forms (Iwase et al., 2007). LSD1 histone demethylases contain a SWIRM domain with an amine oxidase domain containing a substrate binding and an FAD-binding part and are only able to demethylate mono- and dimethylated lysines, such as H3K4me1/2 targeted by LSD1 (Stavropoulos et al., 2006;Yang et al., 2006).

It is interesting to note that several members of both families of demethylases displayed an upregulation during cold acclimation before decreasing until 4 h Deacc followed by an increase until 12 h Deacc before returning to the initial NA gene expression levels after 24 h Deacc. This suggests that both demethylase classes are required during cold acclimation and deacclimation. The elevated gene expression of members of both gene families coincides with previous results, where a reduction in H3K9 methylation during short-term cold stress was described (Hu et al., 2012). Research has further shown that *JUMONJI* and *LSD1* genes are linked to regulation of developmental transitions in *A. thaliana.* For example, histone demethylation of the *FLC* gene by *JMJ30* and *JMJ32* controls flowering at warm temperatures (Yang et al., 2010;Gan et al., 2014). Further, *JUMONJI* genes are upregulated under drought stress in peanut (Govind et al., 2009;Shen et al., 2014), in agreement with the upregulation found in ACC samples in this experiment. Lastly, JUMONJI proteins have been associated with changes in the circadian clock (Jones et al., 2010;Lu et al., 2011a). As the samples analyzed here were harvested throughout a time period of 24 h during deacclimation, circadian regulation could have contributed to the expression changes of e.g. *JMJ30* (Lu et al., 2011b). Additional experiments will be necessary to clarify this contribution.

Nevertheless, results for ACC and 24 h Deacc plants are not affected, as these samples were collected at the same time of day as the NA samples. Therefore we conclude that the majority of the investigated histone demethylases of the JUMONJI-type and LSD1 family are upregulated during cold acclimation.

### Pc-G proteins are involved in epigenetic changes during cold acclimation and deacclimation

The Pc-G gene family was discovered in *Drosophila*, where its members encode proteins able to repress the *HOX* genes (Lewis, 1978). The Pc-G gene family encodes a diverse set of proteins with a variety of molecular activities (Sauvageau and Sauvageau, 2010;del Prete et al., 2015). Polycomb proteins act as multiprotein complexes, Polycomb Repressive Complexes 1 (PRC1) and PRC2 in plants and PRC3 in humans (Schwartz and Pirrotta, 2007;Kleinmanns and Schubert, 2014). The PRC1 complex represses genes through mono-ubiquitination of histone H2A and chromatin remodeling (Kleinmanns and Schubert, 2014). Several *PRC1* genes were investigated, such as *RING1a*, *RING1b* as well as *EMBRYONIC FLOWER 1 (EMF1)*. The PRC2 complex mediates the trimethylation of H3K27 (H3K27me3), which results in the repression of transcription through changes in chromatin organization (Kleinmanns and Schubert, 2014;del Prete et al., 2015).

Most genes encoding proteins of the Pc-G displayed an upregulation during cold acclimation followed by either a relatively stable expression or decreased expression during deacclimation. Polycomb proteins have been linked to abiotic and biotic stresses in *A. thaliana.* WD-40 repeat containing protein MSI1 negatively regulates drought-stress responses in *A. thaliana* and a knockout of this gene confers increased drought tolerance (Alexandre et al., 2009).

Further, H3K27me3 levels decrease at the cold-inducible genes *COR15A* and *GOLS3* during cold stress. However upon transfer to ambient temperatures low H3K27me3 were maintained while *COR15A* and *GOLS3* were repressed again. Thus H3K27me3 is not sufficient to inhibit transcription, but the gene activation rather leads to H3K27me3 removal (Kwon et al., 2009). Further research on the regulation of cold-responsive genes by proteins of the Pc-G showed that *EMF1* and *EMF2* repress several cold-regulated genes such as *COR15A* and *CBF1* (Kim et al., 2010b). Similarly, MSI4/FVE was identified in a screen for repressors of *COR15A* and loss of FVE leads to higher freezing tolerance in cold-acclimated plants (Kim et al., 2004). As many cold-inducible genes carry the PRC2 mark H3K27me3 in the warmth, these genetic analyses are consistent with an important function for Pc-G proteins in prevention of precautious expression of cold-inducible genes. Although we found an increase in the expression of most *PRC2* genes during cold acclimation, PRC2 occupancy analyses during cold acclimation will be required to reveal their presence on the cold-inducible genes. Results collected in this work suggest that a higher expression of genes encoding specific PRC2 subunits, such as *WD-40 repeat-containing proteins* (*MSI1-MSI5*), at ACC conditions might have led to an increase or redistribution of H3K27me3, resulting together with other changes in the increased freezing tolerance of *A. thaliana*. Supporting this hypothesis induction of Pc-G genes during cold acclimation has also been reported previously. *Brassica oleracea* displayed induced expression of alfin-like transcription factors, which are interactors of PRC1, during 24 h at 4°C (Kayum et al., 2016) which is consistent with the observed induction of alfin-like genes in our study.

PRC1 and PRC2 proteins are central regulators of vernalization (Song et al., 2012). The only reported Pc-G protein induced by cold is *VERNALIZATION INSENSITIVE 3 (VIN3)* which is only induced after prolonged cold (at least 10 d) (Sung and Amasino, 2004;Kim et al., 2010a). We did not observe this induction as plants only experienced a three days cold period, however, a cold induction of *VRN1*, *CLF, VAL1* and *FIE* was shown in this work. In addition, several vernalization-related genes were regulated by alternative splicing in the cold, including the PRC2 genes *SWN*, *VRN2* and *EMF2* (Table 2). Importantly, all of the alternatively spliced transcripts result in proteins with modified amino acid sequence. Whether these variants have a different function, stability or interaction partners remains to be determined. Wood et al. (2006) revealed that Pc-G proteins are also regulated at the post-translational level as VRN2, CLF, FIE and SWN showed higher protein abundance after prolonged cold, while no changes in steady-state mRNA levels are detected. In conclusion, our and previous work suggest that Pc-G genes are regulated at the transcriptional, post-transcriptional (alternative splicing) and post-translational level. As the PRC2 mediated H3K27me3 appears to be a major mark repressing cold-regulated genes, tight regulation of PRC2 in the cold is important. Whether PRC2 regulation relates to cold acclimation and chilling/freezing tolerance in addition to vernalization remains to be determined.

### The stress responsive *ATX1* gene of the Trx group is cold induced

Trx-G proteins act as antagonists to Pc-G and activate Pc-G target gene transcription by depositing H3K4me3 (del Prete et al., 2015). Consequently, the activity of these genes must be finely tuned by opposing actions of these protein complexes (del Prete et al., 2015). Further Trx-G proteins can also trimethylate H3K36 to activate transcription of target genes and act as an ATP-dependent chromatin remodeling complex, such as proteins containing a SWITCH or Brahma domain (Schuettengruber et al., 2011;del Prete et al., 2015). Trx-G proteins such as *ATX1* (an H3K4 methyltransferase), which plays a role in drought tolerance, have been linked to abiotic and biotic stresses in *A. thaliana* (Ding et al., 2011). This gene, together with *ATX4*, was also highly induced after cold acclimation. A loss of *ATX1* expression results in decreased germination rates, larger stomatal apertures and thus higher transpiration rates, as well as lower drought tolerance (Ding et al., 2011). Further, binding of ATX1 to the gene *9-cis-epoxycarotenoid dioxygenase 3* (*NCED3*), encoding a protein catalyzing the limiting step in ABA synthesis, was observed, and a loss of ATX1 resulted in decreased *NCED3* levels under drought stress (Ding et al., 2011). ATX1 has also been linked to the regulation of the salicylic acid and jasmonic acid pathways via WRKY70, which is a regulator of the plants defense pathway (Alvarez-Venegas et al., 2007).

The gene *ATP-dependent helicase BRAHMA* (*BRM*), encoding an ATP-dependent chromatin remodeling complex, displayed opposite effects to *ATX1* under drought stress, resulting in increased drought tolerance when the gene was non-functional, through repression of *ABA INSENSITIVE5* (Han et al., 2012). While *BRM* expression is not altered by cold, its paralog *CHR12* is induced after cold acclimation. CHR12 is required to arrest growth after the plant is exposed to a stress (drought, heat, salt) (Mlynarova et al., 2007). Thus, its induction may be directly linking growth and stress responses. Three other highly induced genes after cold acclimation, *FAS1*, *FAS2* and *MSI1* encode proteins which form subunits of CHROMATIN ASSEMBLY FACTOR 1 (CAF-1) and are involved in maintaining the cellular organization of the shoot apical meristem (Kaya et al., 2001). The gene *SWI3A* was the only gene of this group which was highly expressed during the first 12 h of deacclimation. It plays an essential role for plant growth and development (Zhou et al., 2003). These results show that there is a finely tuned regulation of several Trx-G genes and chromatin remodelers involved in abiotic stress response and stress release. Regulatory functions of Trx-G during cold stress have not been reported yet. Nevertheless, the involvement of many of these genes in stress responses in plants suggests a possible participation. Furthermore results of this work suggest that genes encoding Trx-G proteins play a role in cold acclimation in *A. thaliana*. Some proteins of Trx-G may also be involved in deacclimation, but further experiments would be required to investigate this.

### DNA methylation may play a role in deacclimation

One group of DNA methylases adds a methyl group onto cytosine residues in higher eukaryotes and have been proposed to control gene expression in plants during development and regulate transposable elements and heterochromatin (Finnegan et al., 1996;Finnegan et al., 1998;Pavlopoulou and Kossida, 2007). Chromomethylases (CMT) are plant-specific and have been linked to symmetric and asymmetric methylation of DNA (Bartee et al., 2001). Changes in the levels of DNA methylation are known to occur during abiotic stresses; however their exact functions and effects are still unclear. For example, it has been observed that a hyper methylation of DNA occurs during salt stress in wheat (Peng and Zhang, 2009). Some of the investigated genes might be involved in deacclimation, e.g. *CMT2* displayed a continued increase in expression up to 12 h Deacc. Furthermore, DNA demethylases such as DEMETER have been previously linked to plant stress responses and *DME* also showed increased expression during deacclimation. In addition, deletion of three DNA demethylases (*DML1, DML2, DML3*) resulted in increased susceptibility to fungal pathogens and therefore a participation in the biotic stress response of plants was proposed (Le et al., 2014).

Interestingly, *DML3* was significantly increased in comparison to ACC at all deacclimation time points and three out of four demethylases (*DML1, DML2, DML3*) were still upregulated at 12 h and/or 24 h Deacc, suggesting that deacclimation is accompanied by a resetting of DNA methylation.

RNAi-related proteins are commonly found in the nucleus and cytoplasm and are well-known to act in post-transcriptional gene silencing in the cytoplasm (Castel and Martienssen, 2013). Dicer and DICER-LIKE (DCL) proteins are key regulators of small RNA biogenesis (RNAi). Only *DCL1* was induced by cold acclimation whereas *DCL3* was significantly up-regulated only after 12 h Deacc. Expression analyses on *DCL* genes in rice showed differential responses comparing drought, cold and salt stress (Liu et al., 2009). However, the cold response of *DCL* genes in rice differed compared to Arabidopsis, which may be due to the fact that rice, in contrast to Arabidopsis, is a chilling sensitive plants.

### Histone variants respond differentially and strongly to cold acclimation and deacclimation

Histone acetylation can occur on 26 potential lysine residues in a nucleosome (Lusser et al., 2001) and is a reversible process. Acetylation can alter the surface of nucleosomes and destabilize it to enhance binding of proteins to transcribed regions (Berger, 2007). Our results indicate an induction of both HDACs and HAC during cold acclimation and deacclimation. HDACs have been previously linked to responses to drought and salt stress, but not cold, in young rice seedlings (Hu et al., 2009). In *Zea mays* HDACs were induced during cold treatment, resulting in deacetylation of histone subunits H3 and H4. In addition, a direct activation of *ZmDREB1* expression by ZmHDACs was suggested (Hu et al., 2011). In Arabidopsis, *HDA6* is involved in cold acclimation through the regulation of cold responsive genes (To et al., 2011;Kim et al., 2012). Similarly, the expression of *HDA3* was highly induced during cold acclimation, and after 2 h Deacc, whereas *HDA6* was not investigated. A similar expression pattern as for *HDA3* was observed for *HD2C*. HD2A, HD2C and HD2D interact with HDA6 and HDA19 in multiprotein complexes (Luo et al., 2017), and HD2C also interacts with BRAHMA, a chromatin remodeler involved in negative regulation of heat responsive genes (Buszewicz et al., 2016). HD2C and the WD-40-repeat containing protein HOS15 interact before binding to the promoters of the cold responsive genes COR15A and COR47. The cold induction of HOS15-mediated chromatin changes promotes HD2C degradation and is correlated with higher histone acetylation levels on the chromatin of COR genes. Additionally HOS15 recruites CBF transcription factors to COR gene promoters (Park et al., 2018). The reported HD2C degradation seems to be contradictory to a higher *HD2C* gene expression under cold conditions, but an analysis of proteins levels will be necessary to compare these studies. Furthermore, the histone acetyltransferase Gcn5 (not investigated in this work) interacts with transcriptional adapter ADA2B, a transcriptional activator of histone acetyltransferases, and T-DNA insertions of *GCN5* lower the induction of *COR* genes during cold acclimation (Stockinger et al., 2001;Vlachonasios et al., 2003). The *ADA2B* gene was also induced during 6 h Deacc in this work, pointing to a possible activation of histone acetyltransferases.

Additionally, the expression of several genes encoding histone variants has been also investigated. Variants of H2A and H3 are incorporated into the chromatin during the interphase of the cell cycle to confer unique properties to the nucleosome (Deal and Henikoff, 2011). Histone variants of the canonical H2A (*HTA2, HTA10, HTA13*), H2A.Z (*HTA8, HTA9* and *HTA11*) and H2A.W (*HTA6, HTA7, HTA12*) were included in the analysis, as well as the histone H1 variants *H1.1*, *H1.2*, *H1.3*, and *HTR1* to *HTR15* from the histone H3 (Jiang and Berger, 2017). The expression of most genes encoding histone variants was induced during cold acclimation and stayed upregulated during deacclimation. Studies on temperature stress have discovered that H2A.Z variants are regulated by a mild increase in ambient temperatures (Kumar and Wigge, 2010). A recent model stresses the importance of the H2A.Z status for the transcriptional regulation of a gene (Asensi-Fabado et al., 2017). Under cold conditions H2A.Z deposition is increased resulting in higher plant sensitivity to changes in temperature. An interesting expression pattern was detected for histone variant *H1.3*, which was not changed after cold acclimation, but was the only gene displaying a significant and stable upregulation after 6 h to 24 h Deacc, indicating that *H1.3* is not induced by cold, but specifically by deacclimation. *H1.3* was also found to be up-regulated after 24 h Deacc compared to ACC conditions in a comparison of three publicly available data sets using microarray and RNA Seq data and was considered to be part of a core set of 25 common up-regulated genes during deacclimation (Vyse et al., 2019). *H1.3* is drought stress-induced in *A. thaliana*, and also responds to ABA treatment (Ascenzi and Gantt, 1997). A model for the action of histone 1 variants suggests that the small and mobile histone H1.3 replaces the canonical histone variants (H1.1 and H1.2) under stress conditions causing hyper methylation, but the influence of this process on transcriptional regulation and physiological responses is not clear yet (Asensi-Fabado et al., 2017). A similar pattern as for *H1.3*, but without induction at 24 h Deacc was found for *HTA12*. An induction of *H1.3* and *HTA12* during later deacclimation time points indicates a possible role of these genes in memorizing a previous stress event.

Overall, this study shows that many chromatin genes are dynamically transcriptionally and post-transcriptionally regulated during the plant cold response and deacclimation. Further work, especially genetic analyses, will be needed to investigate the function of these genes for both processes in more detail. In addition, the modifications set or removed by chromatin enzymes will require a detailed analysis. As hundreds of genes which are stably repressed in non-stress conditions and are targeted by H3K27me3 are activated within minutes after cold exposure, resetting of epigenetic information can be studied during the cold stress response. How this resetting results in memory of stress and/or induces vernalization, is an exciting question to address in the future.

## Acknowledgements

This work was supported through funds for project A3 to DKH and C7 to DS from the Collaborative Research Center 973 funded by the German Research Foundation (DFG, SFB 973). We gratefully acknowledge support to M.P. through the People Program (Marie-Curie Actions) of the European Union’s Seventh Framework Programme (FP7 People: Marie-Curie Actions FP7-MC-IEF) under REA grant agreement 328,713.

---

**Supplemental Figure 1:** Changes in the alternative splicing of vernalization actors during cold exposure and alignments of the translated variants. The expression profiles were obtained using the webservice at https://wyguo.shinyapps.io/atrtd2_profile_app/ (Calixto et al., 2018; Zhang et al., 2017) and the alignments were made using the Needle algorithm using the translated transcripts extracted from AtRTD2. The alignments were plotted using the Sequence Manipulation Suite.

**Supplemental Figure 2:** Expression changes of genes encoding JUMONJI-type and LSD1-type histone demethylases at non-acclimated conditions (NA) and after 2 h, 4 h, 6 h, 12 h and 24 h of deacclimation (Deacc). Gene expression is presented as log2 fold change to cold acclimated conditions (ACC) (Suppl. Table 6). The scale of log2 fold changes ranges from −3 (blue) to 3 (red) with a median of 0 (white). Significance levels are indicated relative to ACC: ***, *P* < 0.001; **, *P* < 0.01; *, *P* < 0.05 *.

**Supplemental Figure 3:** Expression changes of genes encoding proteins of the polycomb group (Pc-G) family at non-acclimated conditions (NA) and after 2 h, 4 h, 6 h, 12 h and 24 h of deacclimation (Deacc). Gene expression is presented as log2 fold change to cold acclimated conditions (ACC) (Suppl. Table 6). The scale of log2 fold changes ranges from −3 (blue) to 3 (red) with a median of 0 (white). Significance levels are indicated relative to ACC: ***, *P* < 0.001; **, *P* < 0.01; *, *P* < 0.05 *.

**Supplemental Figure 4:** Expression changes of genes encoding proteins of the trithorax group (Trx-G) at non-acclimated conditions (NA) and after 2 h, 4 h, 6 h, 12 h and 24 h of deacclimation (Deacc). Gene expression is presented as log2 fold change to cold acclimated conditions (ACC) (Suppl. Table 6). The scale of log2 fold changes ranges from −3 (blue) to 3 (red) with a median of 0 (white). Significance levels are indicated relative to ACC: ***, *P* < 0.001; **, *P* < 0.01; *, *P* < 0.05 *.

**Supplemental Figure 5:** Expression changes of genes encoding proteins acting in chromosome-nuclear envelope (Chr-NE) interactions, RNA interference and methylation at non-acclimated conditions (NA) and after 2 h, 4 h, 6 h, 12 h and 24 h of deacclimation (Deacc). Gene expression is presented as log2 fold change to cold acclimated conditions (ACC) (Suppl. Table 6). The scale of log2 fold changes ranges from −3 (blue) to 3 (red) with a median of 0 (white). Significance levels are indicated relative to ACC: ***, *P* < 0.001; **, *P* < 0.01; *, *P* < 0.05 *.

**Supplemental Figure 6:** Expression changes of genes encoding histone acetyltransferases (HAC), deacetylases (HDAC) or histone variants at non-acclimated conditions (NA) and after 2 h, 4 h, 6 h, 12 h and 24 h of deacclimation (Deacc). Gene expression is presented as log2 fold change to cold acclimated conditions (ACC) (Suppl. Table 6). The scale of log2 fold changes ranges from −3 (blue) to 3 (red) with a median of 0 (white). Significance levels are indicated relative to ACC: ***, *P* < 0.001; **, *P* < 0.01; *, *P* < 0.05 *.

**Supplemental Table 1:** Primer sequences of 135 genes encoding epigenetic regulators.

**Supplemental Table 2:** Primer sequences of housekeeping genes as well as primer pairs for quality control of the RNA extraction and cDNA synthesis.

**Supplemental Table 3:** Selection of genes induced or repressed late (3 d) or early (overlap of 3 h and 6 h) in the cold, based on published data (Calixto et al., 2018). Asterisks indicate **** = *P* < 0.0001; *** = *P* < 0.001; ** = *P* < 0.01; * = *P* < 0.05.

**Supplemental Table 4:** List of genes in the different sections of the Venn Diagram from Fig. 2, representing the overlap between genes differentially expressed (DE) or alternatively spliced (DAS) during cold exposure, the chromatin modifiers genes described in the ChromDB database (DB) and the genes present on the RT-qPCR platform (RT).

**Supplemental Table 5:** Normalized 2^-ΔCt^ values for all 135 selected genes for non-acclimated (NA) and cold acclimated (ACC) conditions or after 2 h, 4 h, 6 h, 12 h, 24 h deacclimation (Deacc). Values are from three independent experiments.

**Supplemental Table 6:** A) log2 fold change of gene expression of samples from cold acclimated (ACC) conditions or after 2 h, 4 h, 6 h, 12 h, 24 h deacclimation (Deacc) to samples from non-acclimated (NA) conditions for 135 genes encoding epigenetic regulators. B) log2 fold change of gene expression of samples from non-acclimated (NA) conditions or after 2 h, 4 h, 6 h, 12 h, 24 h deacclimation (Deacc) to samples from cold-acclimated (ACC) conditions of the same 135 genes.

## References

Alexandre, C., Möller-Steinbach, Y., Schönrock, N., Gruissem, W., and Hennig, L. (2009). *Arabidopsis* MSI1 is required for negative regulation of the response to drought stress. Molecular Plant 2, 675–687.

Allis, C.D., and Jenuwein, T. (2016). The molecular hallmarks of epigenetic control. Nature Reviews Genetics 17, 487–500.

Alvarez-Venegas, R., Abdallat, A.A., Guo, M., Alfano, J.R., and Avramova, Z. (2007). Epigenetic control of a transcription factor at the cross section of two antagonistic pathways. Epigenetics 2, 106–113.

Alvarez-Venegas, R., Pien, S., Sadder, M., Witmer, X., Grossniklaus, U., and Avramova, Z. (2003). ATX-1, an Arabidopsis homolog of trithorax, activates flower homeotic genes. Current Biology 13, 627–637.

Ascenzi, R., and Gantt, J.S. (1997). A drought-stress-inducible histone gene in *Arabidopsis thaliana* is a member of a distinct class of plant linker histone variants. Plant Molecular Biology 34, 629–641.

Asensi-Fabado, M.-A., Amtmann, A., and Perrella, G. (2017). Plant responses to abiotic stress: The chromatin context of transcriptional regulation. Biochimica et Biophysica Acta (BBA) - Gene Regulatory Mechanisms 1860, 106–122.

Banerjee, A., Wani, S.H., and Roychoudhury, A. (2017). Epigenetic control of plant cold responses. Frontiers in Plant Science 8, 1643–1643.

Bannister, A.J., and Kouzarides, T. (2011). Regulation of chromatin by histone modifications. Cell Research 21, 381–395.

Bartee, L., Malagnac, F., and Bender, J. (2001). *Arabidopsis cmt3* chromomethylase mutations block non-CG methylation and silencing of an endogenous gene. Genes & Development 15, 1753–1758.

Baulcombe, D.C., and Dean, C. (2014). Epigenetic regulation in plant responses to the environment. Cold Spring Harbor Perspectives in Biology 6, a019471.

Berger, S.L. (2007). The complex language of chromatin regulation during transcription. Nature 447, 407–412.

Buszewicz, D., Archacki, R., Palusinski, A., Kotlinski, M., Fogtman, A., Iwanicka-Nowicka, R., Sosnowska, K., Kucinski, J., Pupel, P., Oledzki, J., Dadlez, M., Misicka, A., Jerzmanowski, A., and Koblowska, M.K. (2016). HD2C histone deacetylase and a SWI/SNF chromatin remodelling complex interact and both are involved in mediating the heat stress response in Arabidopsis. Plant, Cell & Environment 39, 2108–2122.

Byun, Y.J., Koo, M.Y., Joo, H.J., Ha-Lee, Y.M., and Lee, D.H. (2014). Comparative analysis of gene expression under cold acclimation, deacclimation and reacclimation in Arabidopsis. Physiologia Plantarum 152, 256–274.

Calixto, C.P.G., Guo, W., James, A.B., Tzioutziou, N.A., Entizne, J.C., Panter, P.E., Knight, H., Nimmo, H.G., Zhang, R., and Brown, J.W.S. (2018). Rapid and dynamic alternative splicing impacts the Arabidopsis cold response transcriptome. Plant Cell 30, 1424–1444.

Carey, N., Marques, C.J., and Reik, W. (2011). DNA demethylases: a new epigenetic frontier in drug discovery. Drug Discovery Today 16, 683–690.

Carles, C.C., and Fletcher, J.C. (2009). The SAND domain protein ULTRAPETALA1 acts as a trithorax group factor to regulate cell fate in plants. Genes & Development 23, 2723–2728.

Castel, S.E., and Martienssen, R.A. (2013). RNA interference (RNAi) in the nucleus: roles for small RNA in transcription, epigenetics and beyond. Nature Reviews Genetics 14, 100–112.

Deal, R.B., and Henikoff, S. (2011). Histone variants and modifications in plant gene regulation. Current Opinion in Plant Biology 14, 116–122.

Del Prete, S., Mikulski, P., Schubert, D., and Gaudin, V. (2015). One, two, three: polycomb proteins hit all dimensions of gene regulation. Genes 6, 520–542.

Ding, Y., Avramova, Z., and Fromm, M. (2011). The *Arabidopsis* trithorax-like factor ATX1 functions in dehydration stress responses via ABA-dependent and ABA-independent pathways. The Plant Journal 66, 735–744.

Finnegan, E.J., Genger, R.K., Peacock, W.J., and Dennis, E.S. (1998). DNA methylation in plants. Annual Review of Plant Physiology and Plant Molecular Biology 49, 223–247.

Finnegan, E.J., Peacock, W.J., and Dennis, E.S. (1996). Reduced DNA methylation in *Arabidopsis thaliana* results in abnormal plant development. Proceedings of the National Academy of Sciences USA 93, 8449–8454.

Friedrich, T., Faivre, L., Baurle, I., and Schubert, D. (2019). Chromatin-based mechanisms of temperature memory in plants. Plant, Cell & Environment 42, 762–770.

Gan, E.-S., Xu, Y., Wong, J.-Y., Goh, J.G., Sun, B., Wee, W.-Y., Huang, J., and Ito, T. (2014). Jumonji demethylases moderate precocious flowering at elevated temperature via regulation of FLC in *Arabidopsis*. Nature Communications 5, 5098.

Gendler, K., Paulsen, T., and Napoli, C. (2008). ChromDB: the chromatin database. Nucleic Acids Research 36, D298–302.

Govind, G., Harshavardhan, V.T., Kalaiarasi, P.J., Iyer, D.R., Muthappa, S.K., Nese, S., and Makarla, U.K. (2009). Identification and functional validation of a unique set of drought induced genes preferentially expressed in response to gradual water stress in peanut. Molecular Genetics and Genomics 281, 591–605.

Han, S.-K., Sang, Y., Rodrigues, A., Wu, M.-F., Rodriguez, P.L., and Wagner, D. (2012). The SWI2/SNF2 chromatin remodeling ATPase BRAHMA represses abscisic acid responses in the absence of the stress stimulus in *Arabidopsis*. The Plant Cell 24, 4892–4906.

He, Y., and Li, Z. (2018). Epigenetic environmental memories in plants: Establishment, maintenance, and reprogramming. Trends in Genetics 34, 856–866.

Hotchkiss, R.D. (1948). The quantitative separation of purines, pyrimidines, and nucleosides by paper chromatography. Journal of Biological Chemistry 175, 315–332.

Hu, Y., Qin, F., Huang, L., Sun, Q., Li, C., Zhao, Y., and Zhou, D.-X. (2009). Rice histone deacetylase genes display specific expression patterns and developmental functions. Biochemical and Biophysical Research Communications 388, 266–271.

Hu, Y., Zhang, L., He, S., Huang, M., Tan, J., Zhao, L., Yan, S., Li, H., Zhou, K., Liang, Y., and Li, L. (2012). Cold stress selectively unsilences tandem repeats in heterochromatin associated with accumulation of H3K9ac. Plant, Cell & Environment 35, 2130–2142.

Hu, Y., Zhang, L., Zhao, L., Li, J., He, S., Zhou, K., Yang, F., Huang, M., Jiang, L., and Li, L. (2011). Trichostatin A selectively suppresses the cold-induced transcription of the ZmDREB1 gene in maize. PLOS ONE 6, e22132.

Iwase, S., Lan, F., Bayliss, P., De La Torre-Ubieta, L., Huarte, M., Qi, H.H., Whetstine, J. R., Bonni, A., Roberts, T.M., and Shi, Y. (2007). The X-linked mental retardation gene *SMCX/JARID1C* defines a family of histone H3 lysine 4 demethylases. Cell 128, 1077–1088.

Jiang, D., and Berger, F. (2017). Histone variants in plant transcriptional regulation. Biochimica et Biophysica Acta - Gene Regulatory Mechanisms 1860, 123–130.

Jin, B., Li, Y., and Robertson, K.D. (2011). DNA methylation: superior or subordinate in the epigenetic hierarchy? Genes & Cancer 2, 607–617.

Jones, M.A., Covington, M.F., Ditacchio, L., Vollmers, C., Panda, S., and Harmer, S.L. (2010). Jumonji domain protein JMJD5 functions in both the plant and human circadian systems. Proceedings of the National Academy of Sciences USA 107, 21623–21628.

Kawashima, T., Lorković, Z.J., Nishihama, R., Ishizaki, K., Axelsson, E., Yelagandula, R., Kohchi, T., and Berger, F. (2015). Diversification of histone H2A variants during plant evolution. Trends in Plant Science 20, 419–425.

Kaya, H., Shibahara, K.I., Taoka, K.I., Iwabuchi, M., Stillman, B., and Araki, T. (2001). FASCIATA genes for chromatin assembly factor-1 in Arabidopsis maintain the cellular organization of apical meristems. Cell 104, 131–142.

Kayum, M.A., Park, J.-I., Ahmed, N.U., Saha, G., Chung, M.-Y., Kang, J.-G., and Nou, I.-S. (2016). Alfin-like transcription factor family: characterization and expression profiling against stresses in *Brassica oleracea*. Acta Physiologiae Plantarum 38, 127.

Kim, D.-H., Zografos, B.R., and Sung, S. (2010a). Vernalization-mediated *VIN3* induction overcomes the LIKE-HETEROCHROMATIN PROTEIN1/POLYCOMB REPRESSION COMPLEX2-mediated epigenetic repression. Plant Physiology 154, 949–957.

Kim, H.J., Hyun, Y., Park, J.Y., Park, M.J., Park, M.K., Kim, M.D., Kim, H.J., Lee, M.H., Moon, J., Lee, I., and Kim, J. (2004). A genetic link between cold responses and flowering time through FVE in Arabidopsis thaliana. Nature Genetics 36, 167–171.

Kim, J.M., Sasaki, T., Ueda, M., Sako, K., and Seki, M. (2015). Chromatin changes in response to drought, salinity, heat, and cold stresses in plants. Frontiers in Plant Science 6, 114.

Kim, J.M., To, T.K., and Seki, M. (2012). An epigenetic integrator: new insights into genome regulation, environmental stress responses and developmental controls by histone deacetylase 6. Plant and Cell Physiology 53, 794–800.

Kim, S.Y., Zhu, T., and Sung, Z.R. (2010b). Epigenetic regulation of gene programs by *EMF1* and *EMF2* in *Arabidopsis*. Plant Physiology 152, 516–528.

Kleinmanns, J.A., and Schubert, D. (2014). Polycomb and Trithorax group protein-mediated control of stress responses in plants. Biological Chemistry 395, 1291–1300.

Kotliński, M., Rutowicz, K., Kniżewski, Ł., Palusiński, A., Olędzki, J., Fogtman, A., Rubel, T., Koblowska, M., Dadlez, M., Ginalski, K., and Jerzmanowski, A. (2016). Histone H1 variants in *Arabidopsis* are subject to numerous post-translational modifications, both conserved and previously unknown in histones, suggesting complex functions of H1 in Plants. PLoS ONE 11, e0147908.

Kumar, S.V., and Wigge, P.A. (2010). H2A.Z-containing nucleosomes mediate the thermosensory response in *Arabidopsis*. Cell 140, 136–147.

Kwon, C.S., Lee, D., Choi, G., and Chung, W.-I. (2009). Histone occupancy-dependent and - independent removal of H3K27 trimethylation at cold-responsive genes in *Arabidopsis*. The Plant Journal 60, 112–121.

Lafos, M., Kroll, P., Hohenstatt, M.L., Thorpe, F.L., Clarenz, O., and Schubert, D. (2011). Dynamic regulation of H3K27 trimethylation during Arabidopsis differentiation. PLoS Genet 7, e1002040.

Law, J.A., and Jacobsen, S.E. (2010). Establishing, maintaining and modifying DNA methylation patterns in plants and animals. Nature Reviews Genetics 11, 204–220.

Le, M.Q., Pagter, M., and Hincha, D.K. (2015). Global changes in gene expression, assayed by microarray hybridization and quantitative RT-PCR, during acclimation of three *Arabidopsis thaliana* accessions to sub-zero temperatures after cold acclimation. Plant Molecular Biology 87, 1–15.

Le, T.-N., Schumann, U., Smith, N.A., Tiwari, S., Au, P.C.K., Zhu, Q.-H., Taylor, J.M., Kazan, K., Llewellyn, D.J., Zhang, R., Dennis, E.S., and Wang, M.-B. (2014). DNA demethylases target promoter transposable elements to positively regulate stress responsive genes in *Arabidopsis*. Genome Biology 15, 458.

Lewis, E.B. (1978). A gene complex controlling segmentation in *Drosophila*. Nature 276, 565–570.

Liu, Q., Feng, Y., and Zhu, Z. (2009). Dicer-like (DCL) proteins in plants. Functional & Integrative Genomics 9, 277–286.

Lu, F., Cui, X., Zhang, S., Jenuwein, T., and Cao, X. (2011a). Arabidopsis REF6 is a histone H3 lysine 27 demethylase. Nature Genetics 43, 715.

Lu, S.X., Knowles, S.M., Webb, C.J., Celaya, R.B., Cha, C., Siu, J.P., and Tobin, E.M. (2011b). The Jumonji C domain-containing protein JMJ30 regulates period length in the *Arabidopsis* circadian clock. Plant Physiology 155, 906–915.

Luger, K., Mader, A.W., Richmond, R.K., Sargent, D.F., and Richmond, T.J. (1997). Crystal structure of the nucleosome core particle at 2.8 Å resolution. Nature 389, 251–260.

Luo, M., Cheng, K., Xu, Y., Yang, S., and Wu, K. (2017). Plant responses to abiotic stress regulated by histone deacetylases. Frontiers in Plant Science 8, 2147.

Lusser, A., Kölle, D., and Loidl, P. (2001). Histone acetylation: lessons from the plant kingdom. Trends in Plant Science 6, 59–65.

Madeira, F., Park, Y.M., Lee, J., Buso, N., Gur, T., Madhusoodanan, N., Basutkar, P., Tivey, A.R.N., Potter, S.C., Finn, R.D., and Lopez, R. (2019). The EMBL-EBI search and sequence analysis tools APIs in 2019. Nucleic Acids Research 47, W636–w641.

Mlynarova, L., Nap, J.P., and Bisseling, T. (2007). The SWI/SNF chromatin-remodeling gene AtCHR12 mediates temporary growth arrest in *Arabidopsis thaliana* upon perceiving environmental stress. Plant Journal 51, 874–885.

Mosammaparast, N., and Shi, Y. (2010). Reversal of histone methylation: biochemical and molecular mechanisms of histone demethylases. Annual Review of Biochemistry 79, 155–179.

Ng, S.S., Yue, W.W., Oppermann, U., and Klose, R.J. (2009). Dynamic protein methylation in chromatin biology. Cellular and Molecular Life Sciences 66, 407–422.

Pagter, M., Alpers, J., Erban, A., Kopka, J., Zuther, E., and Hincha, D.K. (2017). Rapid transcriptional and metabolic regulation of the deacclimation process in cold acclimated *Arabidopsis thaliana*. BMC Genomics 18, 731.

Park, J., Lim, C.J., Shen, M., Park, H.J., Cha, J.-Y., Iniesto, E., Rubio, V., Mengiste, T., Zhu, J.-K., Bressan, R.A., Lee, S.Y., Lee, B.-H., Jin, J.B., Pardo, J.M., Kim, W.-Y., and Yun, D.-J. (2018). Epigenetic switch from repressive to permissive chromatin in response to cold stress. Proceedings of the National Academy of Sciences 115, E5400–E5409.

Pavlopoulou, A., and Kossida, S. (2007). Plant cytosine-5 DNA methyltransferases: structure, function, and molecular evolution. Genomics 90, 530–541.

Peng, H., and Zhang, J. (2009). Plant genomic DNA methylation in response to stresses: Potential applications and challenges in plant breeding. Progress in Natural Science 19, 1037–1045.

Penterman, J., Zilberman, D., Huh, J.H., Ballinger, T., Henikoff, S., and Fischer, R.L. (2007). DNA demethylation in the *Arabidopsis* genome. Proceedings of the National Academy of Sciences USA 104, 6752–6757.

R Core Team (2013). “R: A language and environment for statistical computing.”. R-3.4.3 ed. (Vienna, Austria: R Foundation for Statistical Computing).

Razin, A., and Riggs, A. (1980). DNA methylation and gene function. Science 210, 604–610.

Roy, D., Paul, A., Roy, A., Ghosh, R., Ganguly, P., and Chaudhuri, S. (2014). Differential acetylation of histone H3 at the regulatory region of OsDREB1b promoter facilitates chromatin remodelling and transcription activation during cold stress. PLOS ONE 9, e100343.

Rstudio Team (2016). “RStudio: Integrated Development Environment for R”. 1.1.423 ed. (Boston, MA).

Rutowicz, K., Puzio, M., Halibart-Puzio, J., Lirski, M., Kotlinski, M., Kroten, M.A., Knizewski, L., Lange, B., Muszewska, A., Sniegowska-Swierk, K., Koscielniak, J., Iwanicka-Nowicka, R., Buza, K., Janowiak, F., Zmuda, K., Joesaar, I., Laskowska-Kaszub, K., Fogtman, A., Kollist, H., Zielenkiewicz, P., Tiuryn, J., Siedlecki, P., Swiezewski, S., Ginalski, K., Koblowska, M., Archacki, R., Wilczynski, B., Rapacz, M., and Jerzmanowski, A. (2015). A specialized histone H1 variant is required for adaptive responses to complex abiotic stress and related DNA methylation in Arabidopsis. Plant Physiology 169, 2080–2101.

Sauvageau, M., and Sauvageau, G. (2010). Polycomb group proteins: multi-faceted regulators of somatic stem cells and cancer. Cell Stem Cell 7, 299–313.

Schubert, D., Primavesi, L., Bishopp, A., Roberts, G., Doonan, J., Jenuwein, T., and Goodrich, J. (2006). Silencing by plant Polycomb-group genes requires dispersed trimethylation of histone H3 at lysine 27. Embo Journal 25, 4638–4649.

Schuettengruber, B., Martinez, A.-M., Iovino, N., and Cavalli, G. (2011). Trithorax group proteins: switching genes on and keeping them active. Nature Reviews Molecular Cell Biology 12, 799–814.

Schwartz, Y.B., and Pirrotta, V. (2007). Polycomb silencing mechanisms and the management of genomic programmes. Nature Reviews Genetics 8, 9–22.

Shen, Y., Conde E Silva, N., Audonnet, L., Servet, C., Wei, W., and Zhou, D.-X. (2014). Over-expression of histone H3K4 demethylase gene JMJ15 enhances salt tolerance in *Arabidopsis*. Frontiers in Plant Science 5, 290.

Shi, Y., Lan, F., Matson, C., Mulligan, P., Whetstine, J.R., Cole, P.A., Casero, R.A., and Shi, Y. (2004). Histone demethylation mediated by the nuclear amine oxidase homolog LSD1. Cell 119, 941–953.

Song, J., Angel, A., Howard, M., and Dean, C. (2012). Vernalization - a cold-induced epigenetic switch. Journal of Cell Science 125, 3723–3731.

Spiker, S. (1982). Histone variants in plants. Evidence for primary structure variants differing in molecular weight. Journal of Biological Chemistry 257, 14250–14255.

Stavropoulos, P., Blobel, G., and Hoelz, A. (2006). Crystal structure and mechanism of human lysine-specific demethylase-1. Nature Structural & Molecular Biology 13, 626–632.

Stockinger, E.J., Mao, Y., Regier, M.K., Triezenberg, S.J., and Thomashow, M.F. (2001). Transcriptional adaptor and histone acetyltransferase proteins in *Arabidopsis* and their interactions with CBF1, a transcriptional activator involved in cold-regulated gene expression. Nucleic Acids Research 29, 1524–1533.

Stothard, P. (2000). The sequence manipulation suite: JavaScript programs for analyzing and formatting protein and DNA sequences. Biotechniques 28, 1102, 1104.

Stroud, H., Otero, S., Desvoyes, B., Ramírez-Parra, E., Jacobsen, S.E., and Gutierrez, C. (2012). Genome-wide analysis of histone H3.1 and H3.3 variants in *Arabidopsis thaliana*. Proceedings of the National Academy of Sciences USA 109, 5370–5375.

Sung, S., and Amasino, R.M. (2004). Vernalization in *Arabidopsis thaliana* is mediated by the PHD finger protein VIN3. Nature 427, 159–164.

To, T.K., Nakaminami, K., Kim, J.-M., Morosawa, T., Ishida, J., Tanaka, M., Yokoyama, S., Shinozaki, K., and Seki, M. (2011). Arabidopsis HDA6 is required for freezing tolerance. Biochemical and Biophysical Research Communications 406, 414–419.

Vlachonasios, K.E., Thomashow, M.F., and Triezenberg, S.J. (2003). Disruption mutations of *ADA2b* and *GCN5* transcriptional adaptor genes dramatically affect *Arabidopsis* growth, development, and gene expression. The Plant Cell 15, 626–638.

Vyse, K., Pagter, M., Zuther, E., and Hincha, D.K. (2019). Deacclimation after cold acclimation - a crucial, but widely neglected part of plant winter survival. Journal of Experimental Botany, DOI: 10.1093/jxb/erz229.

Weber, C.M., and Henikoff, S. (2014). Histone variants: dynamic punctuation in transcription. Genes & Development 28, 672–682.

Widom, J. (1989). Toward a unified model of chromatin folding. Annual Review of Biophysics and Biophysical Chemistry 18, 365–395.

Wood, C.C., Robertson, M., Tanner, G., Peacock, W.J., Dennis, E.S., and Helliwell, C.A. (2006). The *Arabidopsis thaliana* vernalization response requires a polycomb-like protein complex that also includes VERNALIZATION INSENSITIVE 3. Proceedings of the National Academy of Sciences USA 103, 14631–14636.

Yang, M., Gocke, C.B., Luo, X., Borek, D., Tomchick, D.R., Machius, M., Otwinowski, Z., and Yu, H. (2006). Structural basis for CoREST-dependent demethylation of nucleosomes by the human LSD1 histone demethylase. Molecular Cell 23, 377–387.

Yang, W., Jiang, D., Jiang, J., and He, Y. (2010). A plant-specific histone H3 lysine 4 demethylase represses the floral transition in *Arabidopsis*. The Plant Journal 62, 663–673.

Yelagandula, R., Stroud, H., Holec, S., Zhou, K., Feng, S., Zhong, X., Muthurajan, Uma m., Nie, X., Kawashima, T., Groth, M., Luger, K., Jacobsen, Steven e., and Berger, F. (2014). The histone variant H2A.W defines heterochromatin and promotes chromatin condensation in *Arabidopsis*. Cell 158, 98–109.

Zeng, Z., Zhang, W., Marand, A.P., Zhu, B., Buell, C.R., and Jiang, J. (2019). Cold stress induces enhanced chromatin accessibility and bivalent histone modifications H3K4me3 and H3K27me3 of active genes in potato. Genome Biology 20, 123.

Zhang, R., Calixto, C.P.G., Marquez, Y., Venhuizen, P., Tzioutziou, N.A., Guo, W., Spensley, M., Entizne, J.C., Lewandowska, D., Ten Have, S., Frei Dit Frey, N., Hirt, H., James, A.B., Nimmo, H.G., Barta, A., Kalyna, M., and Brown, J.W.S. (2017). A high quality Arabidopsis transcriptome for accurate transcript-level analysis of alternative splicing. Nucleic Acids Research 45, 5061–5073.

Zhang, X., Yazaki, J., Sundaresan, A., Cokus, S., Chan, S.W.L., Chen, H., Henderson, I.R., Shinn, P., Pellegrini, M., Jacobsen, S.E., and Ecker, Joseph r. (2006). Genome-wide high-resolution mapping and functional analysis of DNA methylation in *Arabidopsis*. Cell 126, 1189–1201.

Zhang, Y., and Reinberg, D. (2001). Transcription regulation by histone methylation: interplay between different covalent modifications of the core histone tails. Genes & Development 15, 2343–2360.

Zhou, C., Miki, B., and Wu, K. (2003). CHB2, a member of the SWI3 gene family, is a global regulator in Arabidopsis. Plant Molecular Biology 52, 1125–1134.

Zuther, E., Schaarschmidt, S., Fischer, A., Erban, A., Pagter, M., Mubeen, U., Giavalisco, P., Kopka, J., Sprenger, H., and Hincha, D.K. (2019). Molecular signatures associated with increased freezing tolerance due to low temperature memory in Arabidopsis. Plant, Cell & Environment 42, 854–873.

Zuther, E., Schulz, E., Childs, L.H., and Hincha, D.K. (2012). Clinal variation in the non-acclimated and cold-acclimated freezing tolerance of *Arabidopsis thaliana* accessions. Plant, Cell & Environment 35, 1860–1878.

