## Supplemental Data 1 for "Transcriptional and post-transcriptional regulation and transcriptional memory of chromatin regulators in response to low temperature"

# SR45 – AT1G16610

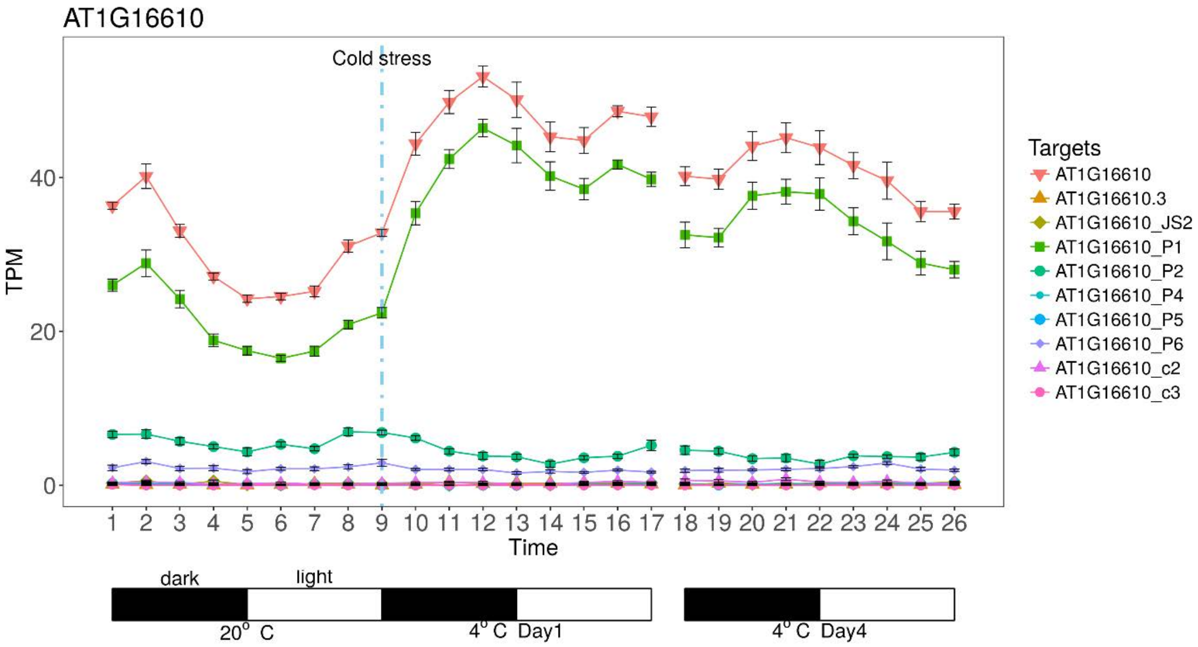


```
AT1G16610_P1   MAKPSRGRRSPSVSGSSSRSSSRSRSGSSPSRSISRSRSRSRSLSSSSSP   50
AT1G16610_P2   MAKPSRGRRSPSVSGSSSRSSSRSRSGSSPSRSISRSRSRSRSLSSSSSP   50

AT1G16610_P1   SRSVSSGSRSPPRRGKSPAGPARRGRSPPPPPSKGASSPSKKAVQESLVL   100
AT1G16610_P2   SRSVSSGSRSPPRRGKSPAGPARRGRSPPPPPSKGASSPSKKAVQESLVL   100

AT1G16610_P1   HVDSLSRNVNEAHLKEIFGNFGEVIHVEIAMDRAVNLPRGHGYVEFKARA   150
AT1G16610_P2   HVDSLSRNVNEAHLKEIFGNFGEVIHVEIAMDRAVNLPRGHGYVEFKARA   150

AT1G16610_P1   DAEKAQLYMDGAQIDGKVVKATFTLPPRQKVSSPPKPVSAAPKRDAPKSD   200
AT1G16610_P2   DAEKAQLYMDGAQIDGKVVKATFTLPPRQKVSSPPKPVSAAPKRDAPKSD   200

AT1G16610_P1   NAAADAEKDGGPRRRPRETSPQRKTGLSPRRRSPLPRRGLSPRRRSPDSPH   250
AT1G16610_P2   NAAADAEKDGGPRRRPRER-------LSPRRRSPLPRRGLSPRRRSPDSPH   243

AT1G16610_P1   RRRPGSPIRRRGDTPPRRRPASPSRGRSPSSPPPRRYRSPPRGSPRRIRG   300
AT1G16610_P2   RRRPGSPIRRRGDTPPRRRPASPSRGRSPSSPPPRRYRSPPRGSPRRIRG   293

AT1G16610_P1   SPVRRRSPLPLRRRSPPPRRLRSPPRRSPIRRRSRSPIRRPGRSRSSSIS   350
AT1G16610_P2   SPVRRRSPLPLRRRSPPPRRLRSPPRRSPIRRRSRSPIRRPGRSRSSSIS   343

AT1G16610_P1   PRKGRGPAGRRGRSSSYSSSPSPRRIPRKISRSRSPKRPLRGKRSSSNSS   400
AT1G16610_P2   PRKGRGPAGRRGRSSSYSSSPSPRRIPRKISRSRSPKRPLRGKRSSSNSS   393

AT1G16610_P1   SSSSPPPPPPPPRKT                                      414
AT1G16610_P2   SSSSPPPPPPPPRKT                                      407
```

### VRN5 – AT3G24440

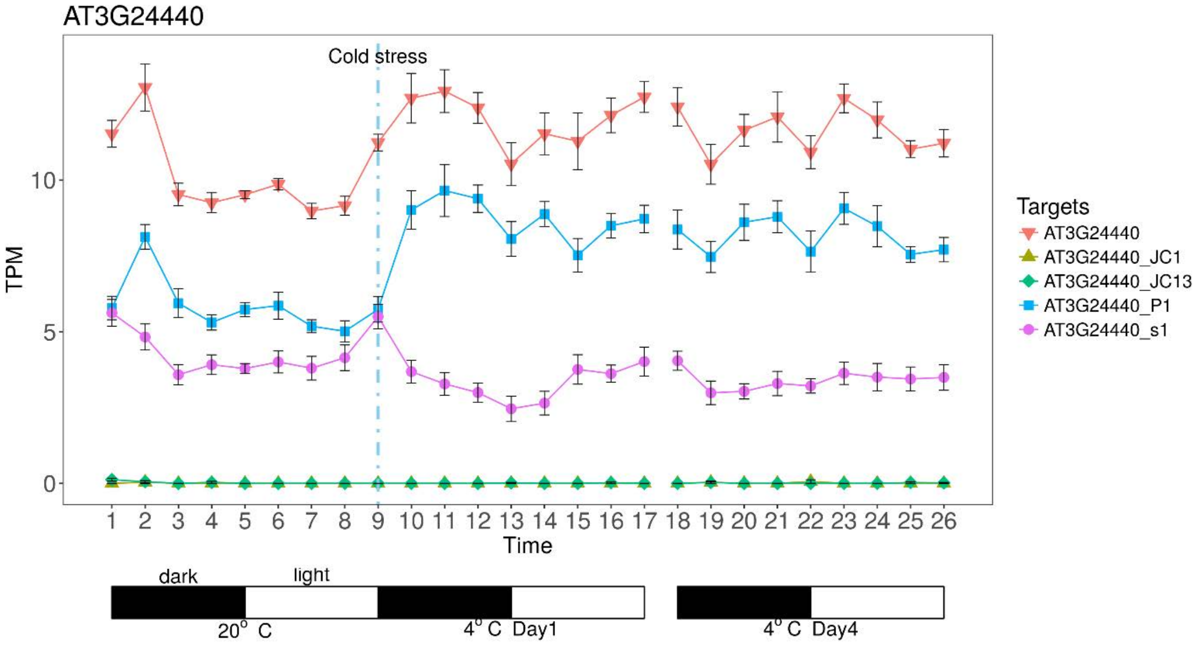


```
AT3G24440_P1  MDSSSTKSKISHSRKTNKKSNKKHESNGKQQQQQDVDGGGGCLRSSWICK   50
AT3G24440_s1  MDSSSTKSKISHSRKTNKKSNKKHESNGKQQQQQDVDGGGGCLRSSWICK   50

AT3G24440_P1  NASCRANVPKEDSFCKRCSCCVCHNFDENKDPSLWLVCEPEKSDDVEFCG  100
AT3G24440_s1  NASCRANVPKEDSFCKRCSCCVCHNFDENKDPSLWLVCEPEKSDDVEFCG  100

AT3G24440_P1  LSCHIECAFREVKVGVIALGNLMKLDGCFCCYSCGKVSQILGCWKKQLVA  150
AT3G24440_s1  LSCHIECAFREVKVGVIALGNLMKLDGCFCCYSCGKVSQILGCWKKQLVA  150

AT3G24440_P1  AKEARRRDGLCYRIDLGYRLLNGTSRFSELHEIVRAAKSMLEDEVGPLDG  200
AT3G24440_s1  AKEARRRDGLCYRIDLGYRLLNGTSRFSELHEIVRAAKSMLEDEVGPLDG  200

AT3G24440_P1  PTARTDRGIVSRLPVAANVQELCTSAIKKAGELSANAGRDLVPAACRFHF  250
AT3G24440_s1  PTARTDRGIVSRLPVAANVQELCTSAIKKAGELSANA--DLVPAACRFHF  248

AT3G24440_P1  EDIAPKQVTLRLIELPSAVEYDVKGYKLWYFKKGEMPEDDLFVDCSRTER  300
AT3G24440_s1  EDIAPKQVTLRLIELPSAVEYDVKGYKLWYFKKGEMPEDDLFVDCSRTER  298

AT3G24440_P1  RMVISDLEPCTEYTFRVVSYTEAGIFGHSNAMCFTKSVEILKPVDGKEKR  350
AT3G24440_s1  RMVISDLEPCTEYTFRVVSYTEAGIFGHSNAMCFTKSVEILKPVDGKEKR  348

AT3G24440_P1  TIDLVGNAQPSDREEKSSISSRFQIGQLGKYVQLAEAQEEGLLEAFYNVD  400
AT3G24440_s1  TIDLVGNAQPSDREEKSSISSRFQIGQLGKYVQLAEAQEEGLLEAFYNVD  398

AT3G24440_P1  TEKICEPPEEELPPRRPHGFDLNVVSVPDLNEEFTPPDSSGGEDNGVPLN  450
AT3G24440_s1  TEKICEPPEEELPPRRPHGFDLNVVSVPDLNEEFTPPDSSGGEDNGVPLN  448

AT3G24440_P1  SLAEADGGDHDDNCDDAVSNGRRKNNNDCLVISDGSGDDTGFDFLMTRKR  500
AT3G24440_s1  SLAEADGGDHDDNCDDAVSNGRRKNNNDCLVISDGSGDDTGFDFLMTRKR  498

AT3G24440_P1  KAISDSNDSENHECDSSSIDDTLEKCVKVIRWLEREGHIKTTFRVRFLTW  550
AT3G24440_s1  KAISDSNDSENHECDSSSIDDTLEKCVKVIRWLEREGHIKTTFRVRFLTW  548

AT3G24440_P1  FSMSSTAQEQSVVSTFVQTLEDDPGSLAGQLVDAFTDVVSTKRPNNGVMT  600
AT3G24440_s1  FSMSSTAQEQSVVSTFVQTLEDDPGSLAGQLVDAFTDVVSTKRPNNGVMT  598

AT3G24440_P1  SH                                                  602
AT3G24440_s1  SH                                                  600
```

### SWN - AT4G02020

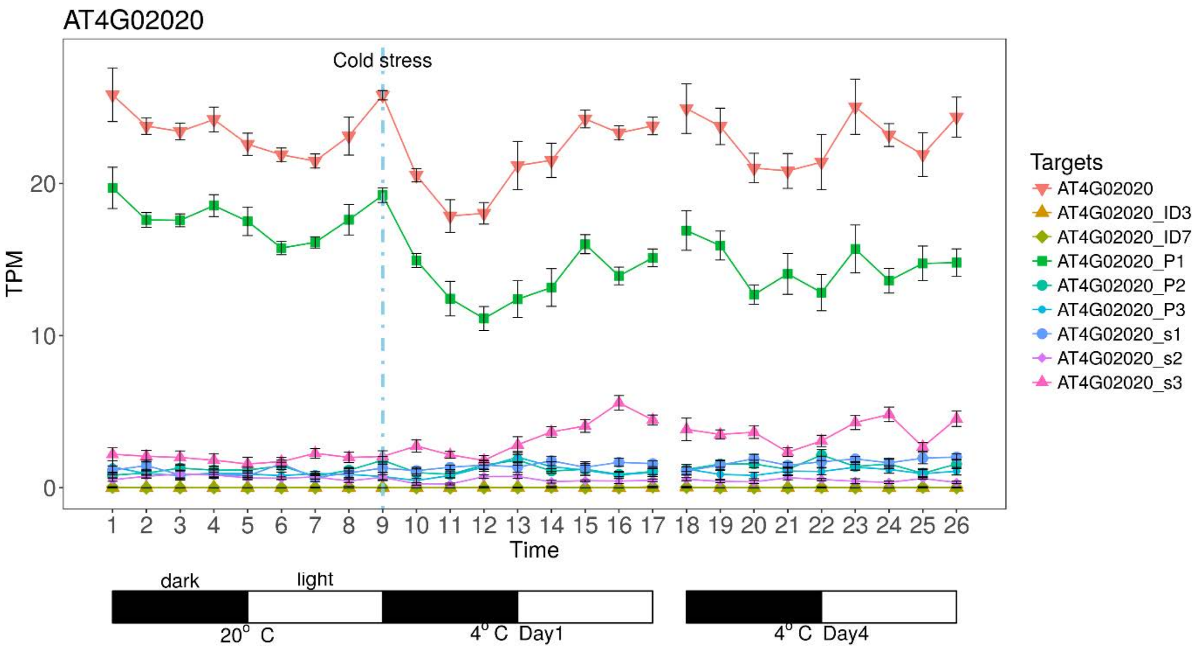

```
AT4G02020_P1  MVTDDSNSSGRIKSHVDDDDDGEEEEDRLEGLENRLSELKRKIQGERVRS  50
AT4G02020_s3  MVTDDSNSSGRI-SHVDDDDDGEEEEDRLEGLENRLSELKRKIQGERVRS  49

AT4G02020_P1  IKEKFEANRKKVDAHVSPFSSAASSRATAEDNGNSNMLSSRMRMPLCKLN  100
AT4G02020_s3  IKEKFEANRKKVDAHVSPFSSAASSRATAEDNGNSNMLSSRMRMPLCKLN  99

AT4G02020_P1  GFSHGVGDRDYVPTKDVISASVKLPIAERIPPYTTWIFLDRNQRMAEDQS  150
AT4G02020_s3  GFSHGVGDRDYVPTKDVISASVKLPIAERIPPYTTWIFLDRNQRMAEDQS  149

AT4G02020_P1  VVGRRQIYYEQHGGETLICSDSEEEPEPEEEKREFSEGEDSIIWLIGQEY  200
AT4G02020_s3  VVGRRQIYYEQHGGETLICSDSEEEPEPEEEKREFSEGEDSIIWLIGQEY  199

AT4G02020_P1  GMGEEVQDALCQLLSVDASDILERYNELKLKDKQNTEEFSNSGFKLGISL  250
AT4G02020_s3  GMGEEVQDALCQLLSVDASDILERYNELKLKDKQNTEEFSNSGFKLGISL  249

AT4G02020_P1  EKGLGAALDSFDNLFCRRCLVFDCRLHGCSQPLISASEKQPYWSDYEGDR  300
AT4G02020_s3  EKGLGAALDSFDNLFCRRCLVFDCRLHGCSQPLISASEKQPYWSDYEGDR  299

AT4G02020_P1  KPCSKHCYLQLKAVREVPETCSNFASKAEEKASEEECSKAVSSDVPHAAA  350
AT4G02020_s3  KPCSKHCYLQLKAVREVPETCSNFASKAEEKASEEECSKAVSSDVPHAAA  349

AT4G02020_P1  SGVSLQVEKTDIGIKNVDSSSGVEQEHGIRGKREVPILKDSNDLPNLSNK  400
AT4G02020_s3  SGVSLQVEKTDIGIKNVDSSSGVEQEHGIRGKREVPILKDSNDLPNLSNK  399

AT4G02020_P1  KQKTAASDTKMSFVNSVPSLDQALDSTKGDQGGTTDNKVNRDSEADAKEV  450
AT4G02020_s3  KQKTAASDTKMSFVNSVPSLDQALDSTKGDQGGTTDNKVNRDSEADAKEV  449

AT4G02020_P1  GEPIPDNSVHDGGSSICQPHHGSGNGAIIIAEMSETSRPSTEWNPIEKDL  500
AT4G02020_s3  GEPIPDNSVHDGGSSICQPHHGSGNGAIIIAEMSETSRPSTEWNPIEKDL  499

AT4G02020_P1  YLKGVEIFGRNSCLIARNLLSGLKTCLDVSNYMRENEVSVFRRSSTPNLL  550
AT4G02020_s3  YLKGVEIFGRNSCLIARNLLSGLKTCLDVSNYMRENEVSVFRRSSTPNLL  549

AT4G02020_P1  LDDGRTDPGNDNDEVPPRTRLFRRKGKTRKLKYSTKSAGHPSVWKRIAGG  600
AT4G02020_s3  LDDGRTDPGNDNDEVPPRTRLFRRKGKTRKLKYSTKSAGHPSVWKRIAGG  599

AT4G02020_P1  KNQSCKQYTPCGCLSMCGKDCPCLTNETCCEKYCGCSKSCKNRFRGCHCA  650
AT4G02020_s3  KNQSCKQYTPCGCLSMCGKDCPCLTNETCCEKYCGCSKSCKNRFRGCHCA  649

AT4G02020_P1  KSQCRSRQCPCFAAGRECDPDVCRNCWVSCGDGSLGEAPRRGEGQCGNMR  700
AT4G02020_s3  KSQCRSRQCPCFAAGRECDPDVCRNCWVSCGDGSLGEAPRRGEGQCGNMR  699

AT4G02020_P1  LLLRQQQRILLGKSDVAGWGAFLKNSVSKNEYLGEYTGELISHHEADKRG  750
AT4G02020_s3  LLLRQQQRILLGKSDVAGWGAFLKNSVSKNEYLGEYTGELISHHEADKRG  749

AT4G02020_P1  KIYDRANSSFLFDLNDQYVLDAQRKGDKLKFANHSAKPNCYAKVMFVAGD  800
AT4G02020_s3  KIYDRANSSFLFDLNDQYVLDAQRKGDKLKFANHSAKPNCYAKVMFVAGD  799

AT4G02020_P1  HRVGIFANERIEASEELFYDYRYGPDQAPVWARKPEGSKKDDSAITHRRA  850
AT4G02020_s3  HRVGIFANERIEASEELFYDYRYGPDQAPVWARKPEGSKKDDSAITHRRA  849

AT4G02020_P1  RKHQSH                                              856
AT4G02020_s3  RKHQSH                                              855
```

### VRN2 - AT4G16845

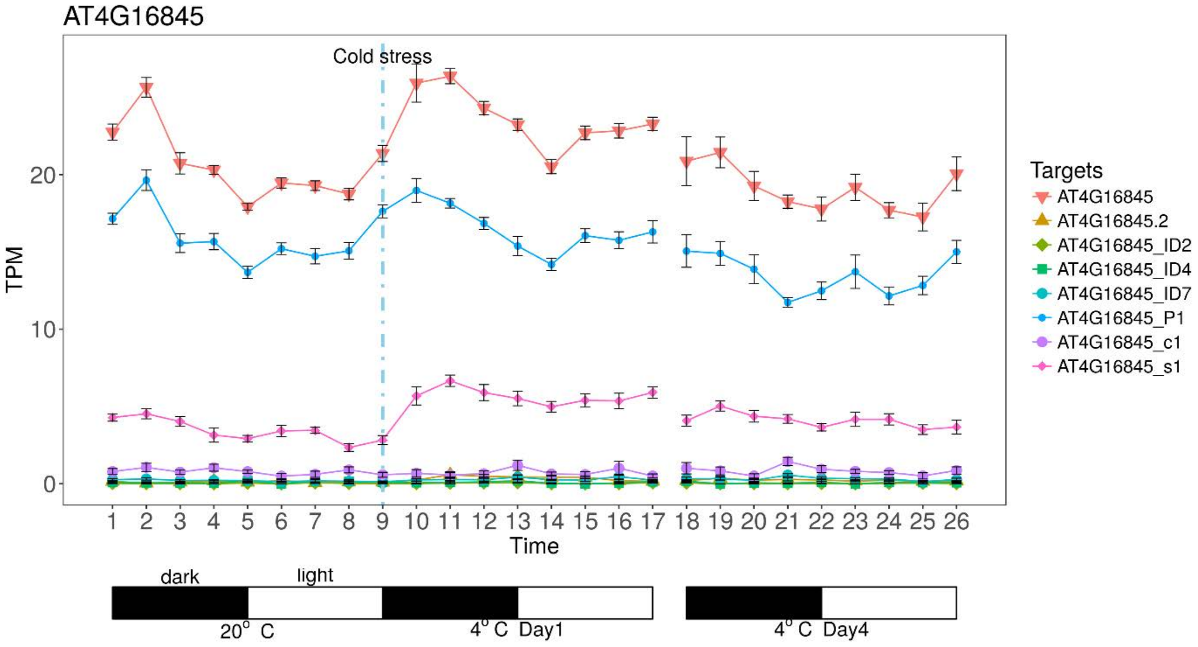


```
AT4G16845_P1  MCRQNCRAKSSPEEVISTDENLLIYCKPVRLYNIFHLRSLGNPSFLPRCL  50
AT4G16845_s1  MCRQNCRAKSSPEEVISTDENLLIYCKPVRLYNIFHLRSLGNPSFLPRCL  50

AT4G16845_P1  NYKIGAKRKRKSRSTGMVVFNYKDCNNTLQRTEVREDCSCPFCSMLCGSF  100
AT4G16845_s1  NYKIGAKRKRKSRSTGMVVFNYKDCNNTLQRTEVREDCSCPFCSMLCGSF  100

AT4G16845_P1  KGLQFHLNSSHDLFEFEFKLLEEYQTVNVSVKLNSFIFEEEGSDDDKFEP  150
AT4G16845_s1  KGLQFHLNSSHDLFEFEFKLLEEYQTVNVSVKLNSFIFEEEGSDDDKFEP  150

AT4G16845_P1  FSLCSKPRKRRQRGGRNNTRRLKVCFLPLDSPSLANGTENGIALLNDGNR  200
AT4G16845_s1  FSLCSKPRKRRQRGGRNNTRRLKVCFLPLDSPSLANGTENGIALLNDGNR  200

AT4G16845_P1  GLGYPEATELAGQFEMTSNIPPAIAHSSLDAGAKVILTTEAVVPATKTRK  250
AT4G16845_s1  GLGYPEATELAGQFEMTSNIPPAIAHSSLDAGAKVILTTEAVVPATKTRK  250

AT4G16845_P1  LSAERSEARSHLLLQKRQFYHSHRVQPMALEQVMSDRDSEDEVDDDVADF  300
AT4G16845_s1  LSAERSEARSHLLLQKRQFYHSHRVQPMALEQVMSDRDSEDEVDDDVADF  300

AT4G16845_P1  EDR--QMLDDFVDVNKDEKQFMHLWNSFVRKQRVIADGHISWACEVFSRF  348
AT4G16845_s1  EDRQLQMLDDFVDVNKDEKQFMHLWNSFVRKQRVIADGHISWACEVFSRF  350

AT4G16845_P1  YEKELHCYSSLFWCWRLFLIKLWNHGLVDSATINNCNTILENCRNTSVTN  398
AT4G16845_s1  YEKELHCYSSLFWCWRLFLIKLWNHGLVDSATINNCNTILENCRNTSVTN  400

AT4G16845_P1  NNNNSVDHPSDSNTNNNNIVDHPNDIKNKNNVDNKDNNSRDK          440
AT4G16845_s1  NNNNSVDHPSDSNTNNNNIVDHPNDIKNKNNVDNKDNNSRDK          442
```

### VEL1/VIL2 – AT4G30200

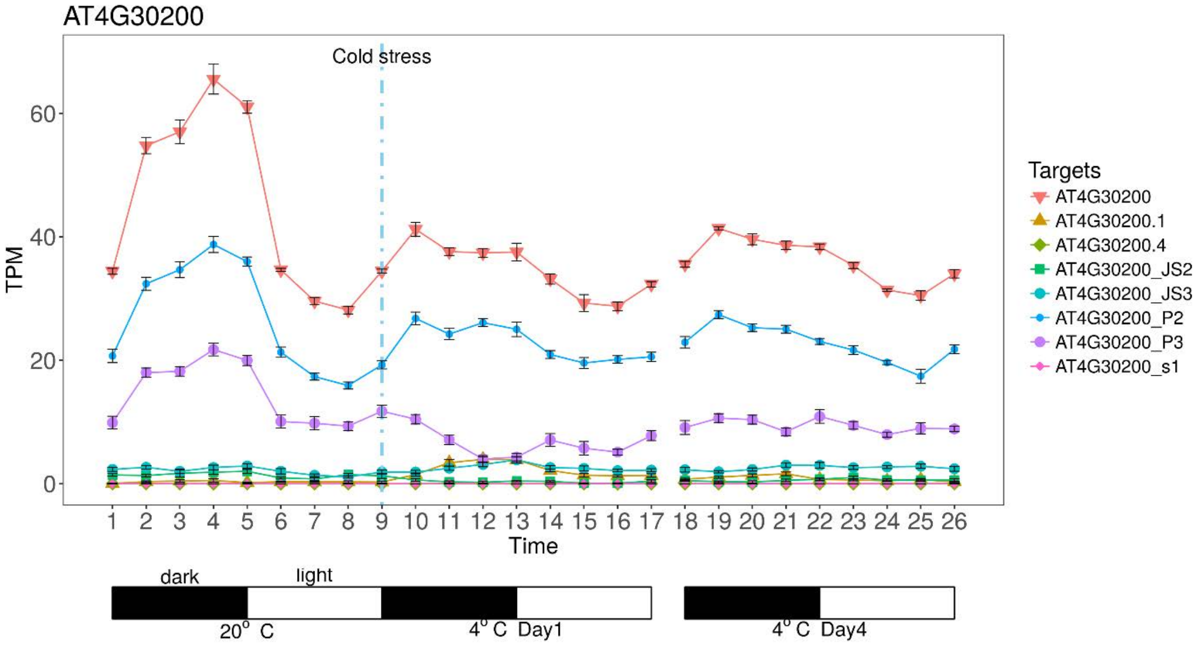

```
AT4G30200_P2  MDSSLDGAAGDSSKCSEMSVDEKRQLVYELSKQSHLAAEVLQAWSRQEIL  50
AT4G30200_P3  MDSSLDGAAGDSSKCSEMSVDEKRQLVYELSKQSHLAAEVLQAWSRQEIL  50

AT4G30200_P2  QILCAEMGKERKYTGLTKVKIIETLLKIVSEKNSGECEGKKRDSDCLPIQ  100
AT4G30200_P3  QILCAEMGKERKYTGLTKVKIIETLLKIVSEKNSGECEGKKRDSDCLPIQ  100

AT4G30200_P2  RNTKRQRKVDNPSRYVIPATNIVTSNNASGSCSSVNTKGESTTIYCKNLA  150
AT4G30200_P3  RNTKRQRKVDNPSRYVIPATNIVTSNNASGSCSSVNTKGESTTIYCKNLA  150

AT4G30200_P2  CRAVLRQEDSFCRRCSCCICRKYDDNKDPSLWLTCSSDPPFEGESCGFSC  200
AT4G30200_P3  CRAVLRQEDSFCRRCSCCICRKYDDNKDPSLWLTCSSDPPFEGESCGFSC  200

AT4G30200_P2  HLECAFNTEKSGLGKDKQSEGCCFYCVSCGKANSLLECWKKQLTIAKETR  250
AT4G30200_P3  HLECAFNTEKSGLGKDKQSEGCCFYCVSCGKANSLLECWKKQLTIAKETR  250

AT4G30200_P2  RVEVLCYRLFLVQKLLKSSTKYRNLCEVVDEAVKTLEADVGPLTGLPMKM  300
AT4G30200_P3  RVEVLCYRLFLVQKLLKSSTKYRNLCEVVDEAVKTLEADVGPLTGLPMKM  300

AT4G30200_P2  GRGIVNRLHSGPDVQKLCSSALESLETIATTPPDVAALPSPRSSKMQQDC  350
AT4G30200_P3  GRGIVNRLHSGPDVQKLCSSALESLETIATTPPDVAALPSPRSSKMQQ--  348

AT4G30200_P2  SYVLSNEISADTATTGSTKIRFEDVNATSLTVVLASNEIPSPPNIVHYSI  400
AT4G30200_P3  ----------DTATTGSTKIRFEDVNATSLTVVLASNEIPSPPNIVHYSI  388

AT4G30200_P2  WHRKVPEKDYPEKSTCTLFIPNTRFVVSGLAPASEYCFKVVSYSGTREMG  450
AT4G30200_P3  WHRKVPEKDYPEKSTCTLFIPNTRFVVSGLAPASEYCFKVVSYSGTREMG  438

AT4G30200_P2  VDEINVLTRSAEEGANCSSAVERSVSPLTNCSTLSSNPSSVEAESNNDYI  500
AT4G30200_P3  VDEINVLTRSAEEGANCSSAVERSVSPLTNCSTLSSNPSSVEAESNNDYI  488

AT4G30200_P2  VPKKPSSKNEDNNSPSVDESAAKRMKRTTDSDIVQIEKDVEQIVLLDDEE  550
AT4G30200_P3  VPKKPSSKNEDNNSPSVDESAAKRMKRTTDSDIVQIEKDVEQIVLLDDEE  538

AT4G30200_P2  QEAVLDKTESETPVVVTTKSLVGNRNSSDASLPITPFRSDEIKNRQARIE  600
AT4G30200_P3  QEAVLDKTESETPVVVTTKSLVGNRNSSDASLPITPFRSDEIKNRQARIE  588

AT4G30200_P2  ISMKDNCNNGDHSANGGTESGLEHCVKIIRQLECSGHIDKNFRQKFLTWY  650
AT4G30200_P3  ISMKDNCNNGDHSANGGTESGLEHCVKIIRQLECSGHIDKNFRQKFLTWY  638

AT4G30200_P2  SLRATSQEIRVVKIFIDTFIDDPMALAEQLIDTFDDRVSIKRSAVGGSGA  700
AT4G30200_P3  SLRATSQEIRVVKIFIDTFIDDPMALAEQLIDTFDDRVSIKRSAVGGSGA  688

AT4G30200_P2  SAVVPSGFCMKLWH                                      714
AT4G30200_P3  SAVVPSGFCMKLWH                                      702
```

### HSL1 – AT4G32010

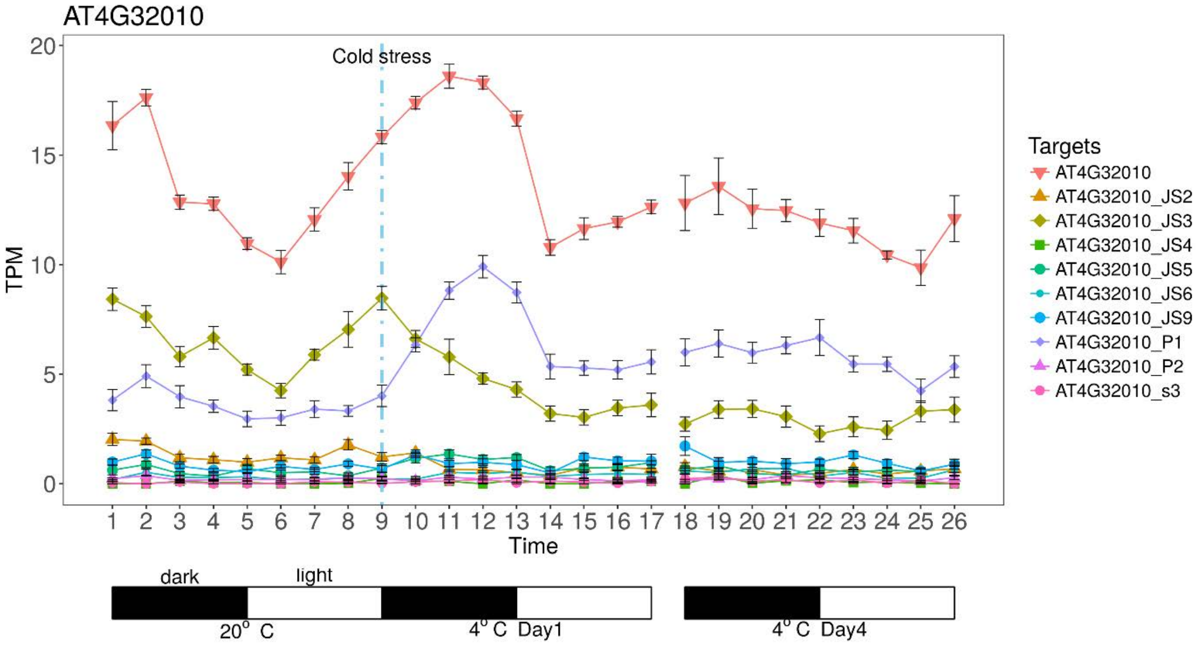

```
AT4G32010_P1   MESIKVCMNALCGAASTSGEWKKGWPMRSGDLASLCDKCGCAYEQSIFCE   50
AT4G32010_JS3  MESIKVCMNALCGAASTSGEWKKGWPMRSGDLASLCDKCGCAYEQSIFCE   50

AT4G32010_P1   VFHAKESGWRECNSCDKRLHCGCIASRFMMELLENGGVTCISCAKKSGLI   100
AT4G32010_JS3  VFHAKESGWRECNSCDKRLHCGCIASRFMMELLENGGVTCISCAKKSGLI   100

AT4G32010_P1   SMNVSHESNGKDFPSFASAEHVGSVLERTNLKHLLHFQRIDPTHSSLQMK   150
AT4G32010_JS3  SMNVSHESNGKDFPSFASAEHVGSVLERTNLKHLLHFQRIDPTHSSLQMK   150

AT4G32010_P1   QEESLLPSSLDALRHKTERKELSAQPNLSISLGPTLMTSPFHDAAVDDRS   200
AT4G32010_JS3  QEESLLPSSLDALRHKTERKELSAQPNLSISLGPTLMTSPFHDAAVDDRS   200

AT4G32010_P1   KTNSIFQLAPRSRQLLPKPANSAPIAAGMEPSGSLVSQIHVARPPPEGRG   250
AT4G32010_JS3  KTNSIFQLAPRSRQLLPKPANSAPIAAGMEPSGSLVSQIHVARPPPEGRG   250

AT4G32010_P1   KTQLLPRYWPRITDQELLQLSGQYPHLSNSKIIPLFEKVLSASDAGRIGR   300
AT4G32010_JS3  KTQLLPRYWPRITDQELLQLSGH----SNSKIIPLFEKVLSASDAGRIGR   296

AT4G32010_P1   LVLPKACAEAYFPPISLPEGLPLKIQDIKGKEWVFQFRFWPNNNSRMYVL   350
AT4G32010_JS3  LVLPKACAEAYFPPISLPEGLPLKIQDIKGKEWVFQFRFWPNNNSRMYVL   346

AT4G32010_P1   EGVTPCIQSMQLQAGDTVTFSRTEPEGKLVMGYRKATNSTATQMFKGSSE   400
AT4G32010_JS3  EGVTPCIQSMQLQAGDTVTFSRTEPEGKLVMGYRKATNSTATQMFKGSSE   396

AT4G32010_P1   PNLNMFSNSLNPGCGDINWSKLEKSEDMAKDNLFLQSSLTSARKRVRNIG   450
AT4G32010_JS3  PNLNMFSNSLNPGCGDINWSKLEKSEDMAKDNLFLQSSLTSARKRVRNIG   446

AT4G32010_P1   TKSKRLLIDSVDVLELKITWEEAQELLRPPQSTKPSIFTLENQDFEEYDE   500
AT4G32010_JS3  TKSKRLLIDSVDVLELKITWEEAQELLRPPQSTKPSIFTLENQDFEEYDE   496

AT4G32010_P1   PPVFGKRTLFVSRQTGEQEQWVQCDACGKWRQLPVDILLPPKWSCSDNLL   550
AT4G32010_JS3  PPVFGKRTLFVSRQTGEQEQWVQCDACGKWRQLPVDILLPPKWSCSDNLL   546

AT4G32010_P1   DPGRSSCSAPDELSPREQDTLVRQSKEFKRRRLASSNEKLNQSQDASALN   600
AT4G32010_JS3  DPGRSSCSAPDELSPREQDTLVRQSKEFKRRRLASSNEKLNQSQDASALN   596

AT4G32010_P1   SLGNAGITTTGEQGEITVAATTKHPRHRAGCSCIVCSQPPSGKGKHKPSC   650
AT4G32010_JS3  SLGNAGITTTGEQGEITVAATTKHPRHRAGCSCIVCSQPPSGKGKHKPSC   646

AT4G32010_P1   TCTVCEAVKRRFRTLMLRKRNKGEAGQASQQAQSQSECRDETEVESIPAV   700
AT4G32010_JS3  TCTVCEAVKRRFRTLMLRKRNKGEAGQASQQAQSQSECRDETEVESIPAV   696

AT4G32010_P1   ELAAGENIDLNSDPGASRVSMMRLLQAAAFPLEAYLKQKAISNTAGEQQS   750
AT4G32010_JS3  ELAAGENIDLNSDPGASRVSMMRLLQAAAFPLEAYLKQKAISNTAGEQQS   746

AT4G32010_P1   SDMVSTEHGSSSAAQETEKDTTNGAHDPVN                       780
AT4G32010_JS3  SDMVSTEHGSSSAAQETEKDTTNGAHDPVN                       776
```

### EMF2 – AT5G51230

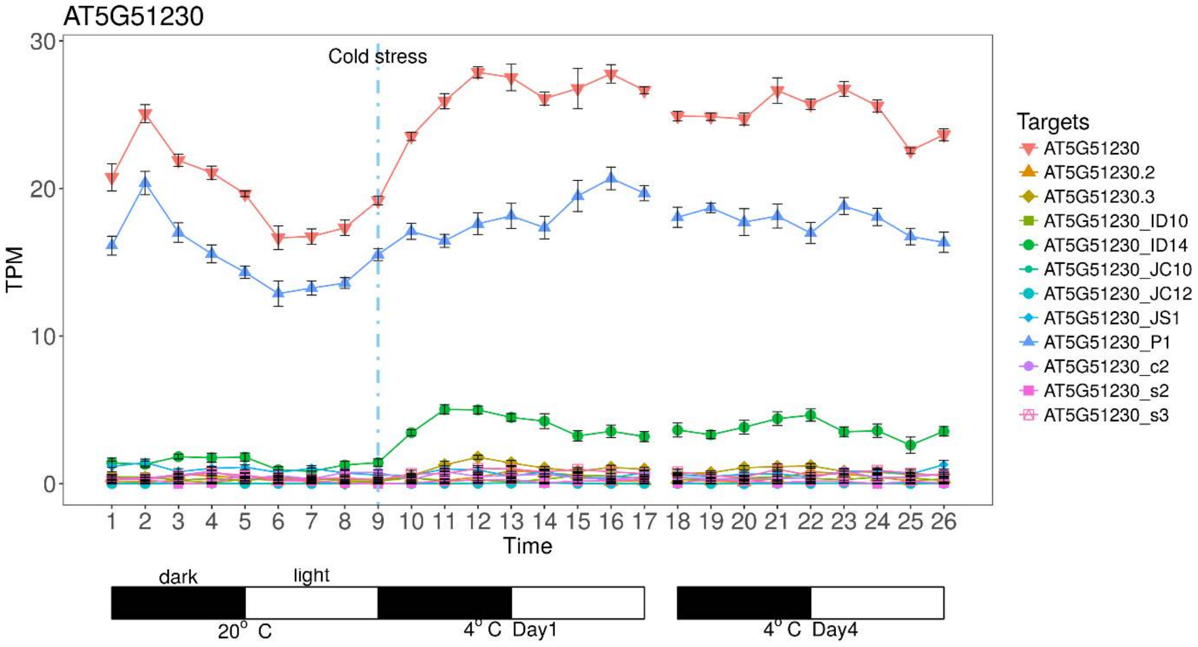


```
AT5G51230_P1    MPGIPLVSRETSSCSRSTEQMCHEDSRLRISEEEEIAAEESLAAYCKPVE  50
AT5G51230_ID14  MPGIPLVSRETSSCSRSTEQMCHEDSRLRISEEEEIAAEESLAAYCKPVE  50

AT5G51230_P1    LYNIIQRRAIRNPLFLQRCLHYKIEAKHKRRIQMTVFLSGAIDAGVQTQK  100
AT5G51230_ID14  LYNIIQRRAIRNPLFLQRCLHYKIEAKHKRRIQMTVFLSGAIDAGVQTQK  100

AT5G51230_P1    LFPLYILLARLVSPKPVAEYSAVYRFSRACILTGGLGVDGVSQAQANFLL  150
AT5G51230_ID14  LFPLYILLARLVSPKPVAEYSAVYRFSRACILTGGLGVDGVSQAQANFLL  150

AT5G51230_P1    PDMNRLALEAKSGSLAILFISFAGAQNSQFGIDSGKIHSGNIGGHCLWSK  200
AT5G51230_ID14  PDMNRLALEAKSGSLAILFISFAGAQNSQFGIDSGKIHSGNIGGHCLWSK  200

AT5G51230_P1    IPLQSLYASWQKSPNMDLGQRVDTVSLVEMQPCFIKLKSMSEEKCVSIQV  250
AT5G51230_ID14  IPLQSLYASWQKSPNMDLGQRVDTVSLVEMQPCFIKLKSMSEEKCVSIQV  250

AT5G51230_P1    PSNPLTSSSPQQVQVTISAEEVGSTEKSPYSSFSYNDISSSSLLQIIRLR  300
AT5G51230_ID14  PSNPLTSSSPQQVQVTISAEEVGSTEKSPYSSFSYNDISSSSLLQIIRLR  300

AT5G51230_P1    TGNVVFNYRYYNNKLQKTEVTEDFSCPFCLVKCASFKGLRYHLPSTHDLL  350
AT5G51230_ID14  TGNVVFNYRYYNNKLQKTEVTEDFSCPFCLVKCASFKGLRYHLPSTHDLL  350

AT5G51230_P1    NFEEFWVTEEFQ-------------AVNVSLKTETMISKVNEDDVDPKQQ  386
AT5G51230_ID14  NFEEFWVC-SFKIQLTCLIFFFYFVGATNLLYLVR---------------  383

AT5G51230_P1    TFFFSSKKFRRRRQKSQVRSSRQGPHLGLGCEVLDKTDDAHSVRSEKSRI  436
AT5G51230_ID14  --------------------------------------------------  383

AT5G51230_P1    PPGKHYERIGGAESGQRVPPGTSPADVQSCGDPDYVQSIAGSTMLQFAKT  486
AT5G51230_ID14  --------------------------------------------------  383

AT5G51230_P1    RKISIERSDLRNRSLLQKRQFFHSHRAQPMALEQVLSDRDSEDEVDDDVA  536
AT5G51230_ID14  --------------------------------------------------  383

AT5G51230_P1    DFEDRRMLDDFVDVTKDEKQMMHMWNSFVRKQRVLADGHIPWACEAFSRL  586
AT5G51230_ID14  --------------------------------------------------  383

AT5G51230_P1    HGPIMVRTPHLIWCWRVFMVKLWNHGLLDARTMNNCNTFLEQLQI       631
AT5G51230_ID14  ---------------------------------------------       383
```

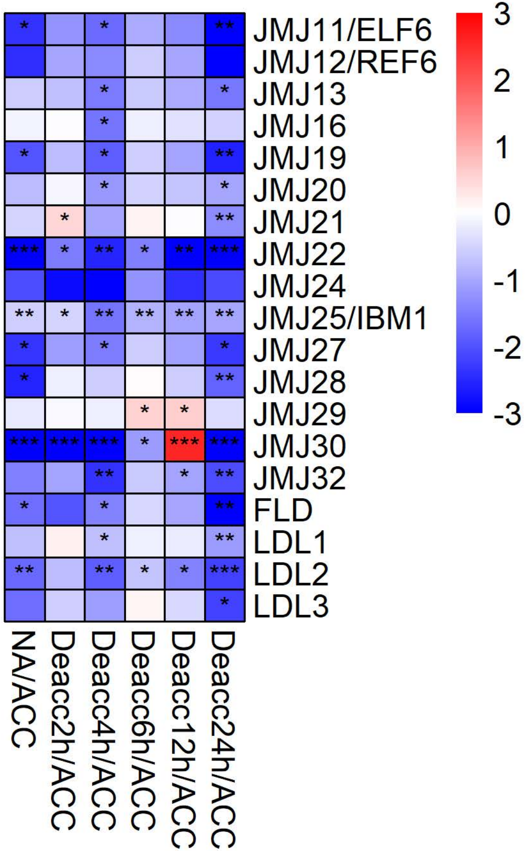

JMJ11/ELF6
JMJ12/REF6
JMJ13
JMJ16
JMJ19
JMJ20
JMJ21
JMJ22
JMJ24
JMJ25/IBM1
JMJ27
JMJ28
JMJ29
JMJ30
JMJ32
FLD
LDL1
LDL2
LDL3
NA/ACC
Deacc2h/ACC
Deacc4h/ACC
Deacc6h/ACC
Deacc12h/ACC
Deacc24h/ACC
3
2
1
0
-1
-2
-3

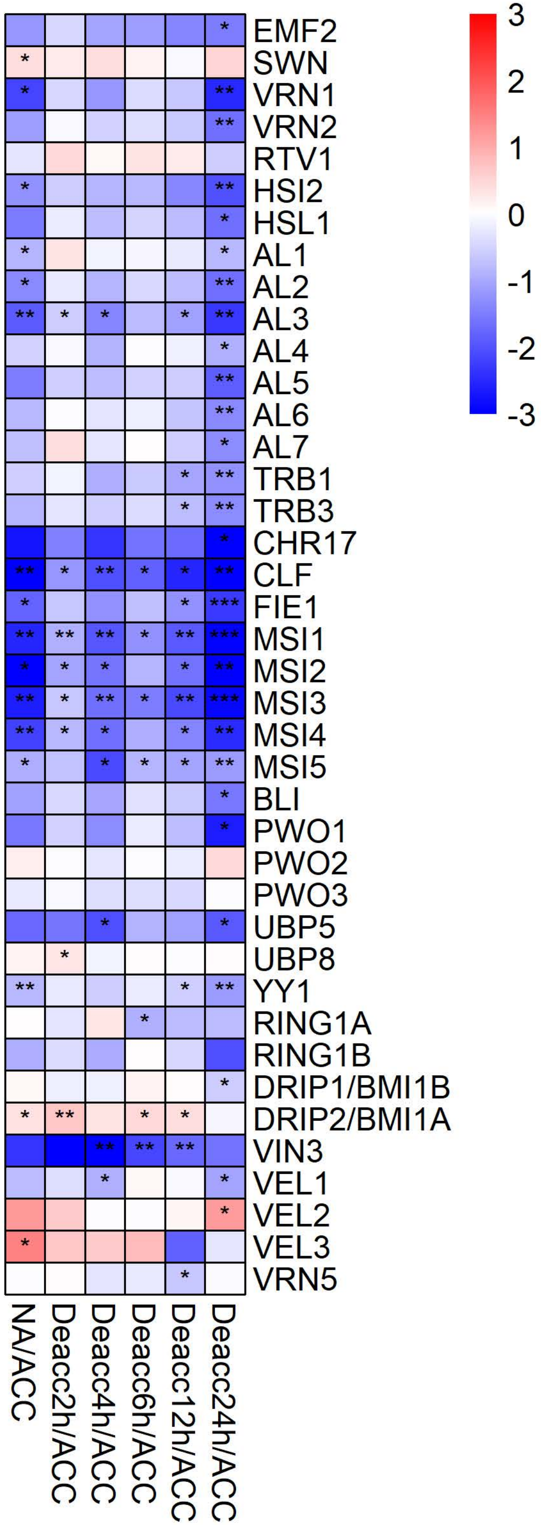
EMF2
SWN
VRN1
VRN2
RTV1
HSI2
HSL1
AL1
AL2
AL3
AL4
AL5
AL6
AL7
TRB1
TRB3
CHR17
CLF
FIE1
MSI1
MSI2
MSI3
MSI4
MSI5
BLI
PWO1
PWO2
PWO3
UBP5
UBP8
YY1
RING1A
RING1B
DRIP1/BMI1B
DRIP2/BMI1A
VIN3
VEL1
VEL2
VEL3
VRN5
NA/ACC
Deacc2h/ACC
Deacc4h/ACC
Deacc6h/ACC
Deacc12h/ACC
Deacc24h/ACC
3
2
1
0
-1
-2
-3

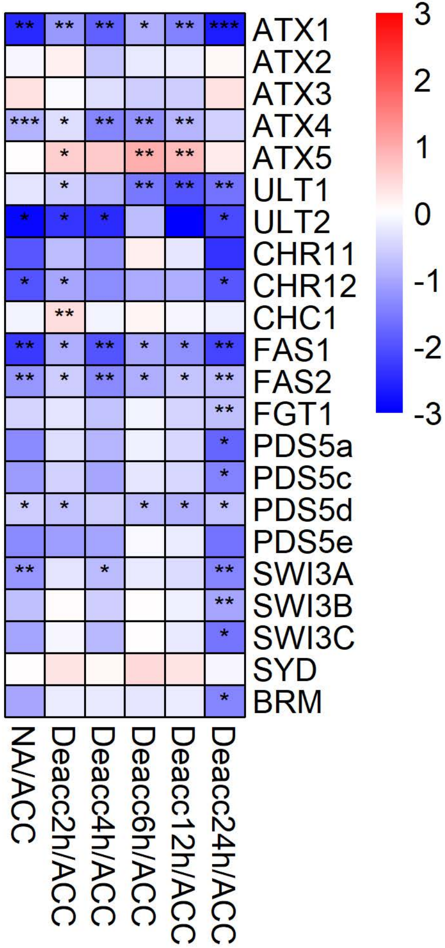

ATX1
ATX2
ATX3
ATX4
ATX5
ULT1
ULT2
CHR11
CHR12
CHC1
FAS1
FAS2
FGT1
PDS5a
PDS5c
PDS5d
PDS5e
SWI3A
SWI3B
SWI3C
SYD
BRM
NA/ACC
Deacc2h/ACC
Deacc4h/ACC
Deacc6h/ACC
Deacc12h/ACC
Deacc24h/ACC
3
2
1
0
-1
-2
-3

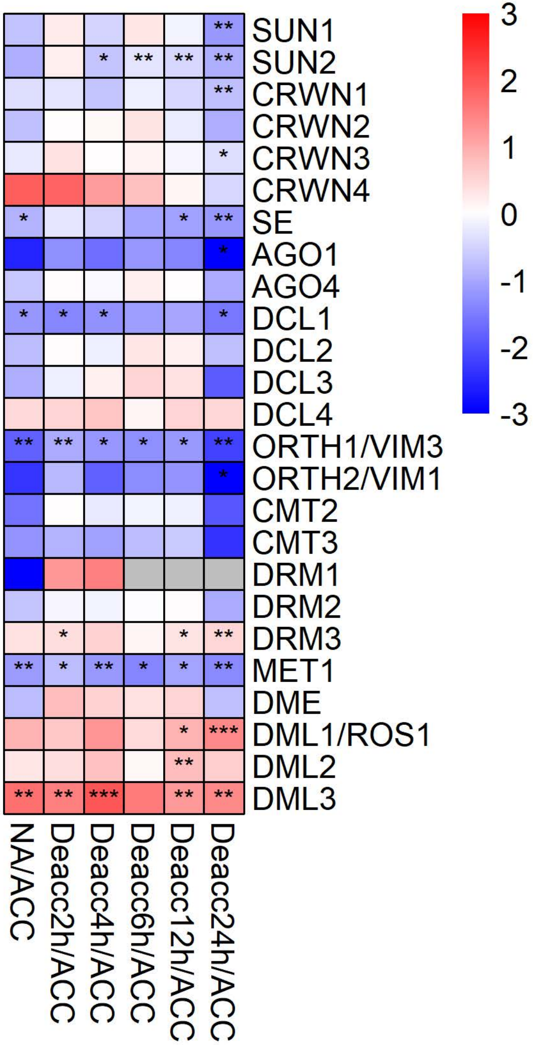
SUN1
SUN2
CRWN1
CRWN2
CRWN3
CRWN4
SE
AGO1
AGO4
DCL1
DCL2
DCL3
DCL4
ORTH1/VIM3
ORTH2/VIM1
CMT2
CMT3
DRM1
DRM2
DRM3
MET1
DME
DML1/ROS1
DML2
DML3
NA/ACC
Deacc2h/ACC
Deacc4h/ACC
Deacc6h/ACC
Deacc12h/ACC
Deacc24h/ACC
3
2
1
0
-1
-2
-3

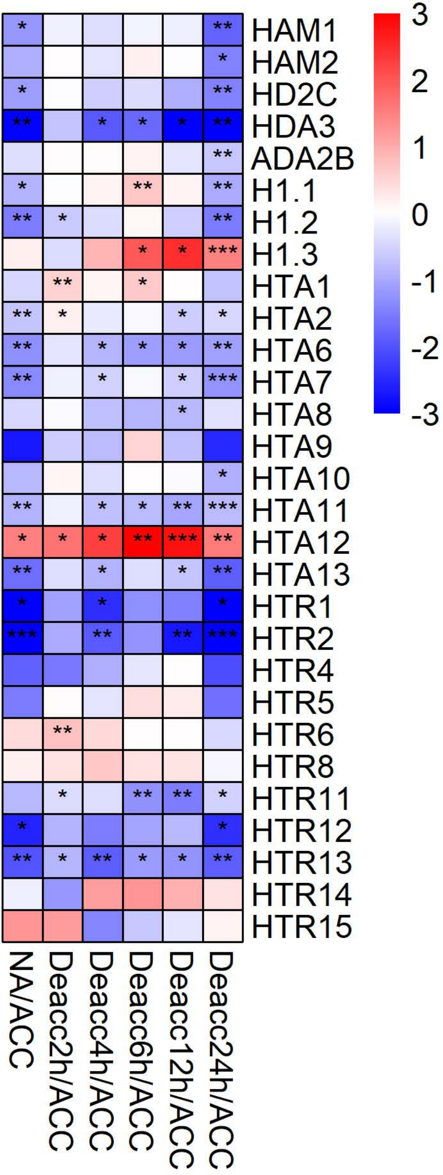

HAM1
HAM2
HD2C
HDA3
ADA2B
H1.1
H1.2
H1.3
HTA1
HTA2
HTA6
HTA7
HTA8
HTA9
HTA10
HTA11
HTA12
HTA13
HTR1
HTR2
HTR4
HTR5
HTR6
HTR8
HTR11
HTR12
HTR13
HTR14
HTR15
NA/ACC
Deacc2h/ACC
Deacc4h/ACC
Deacc6h/ACC
Deacc12h/ACC
Deacc24h/ACC
3
2
1
0
-1
-2
-3
